# Developing and Benchmarking Sulfate and Sulfamate Force Field Parameters for Glycosaminoglycans via Ab Initio Molecular Dynamics Simulations

**DOI:** 10.1101/2024.05.31.596767

**Authors:** Miguel Riopedre-Fernandez, Vojtech Kostal, Tomas Martinek, Hector Martinez-Seara, Denys Biriukov

**Affiliations:** Institute of Organic Chemistry and Biochemistry, Czech Academy of Sciences, Flemingovo nám. 542/2, CZ-16610 Prague 6, Czech Republic; CEITEC – Central European Institute of Technology, Masaryk University, Kamenice 753/5, CZ-62500 Brno, Czech Republic; National Centre for Biomolecular Research, Faculty of Science, Masaryk University, Kamenice 753/5, CZ-62500 Brno, Czech Republic

## Abstract

Glycosaminoglycans (GAGs) are negatively charged polysaccharides found on cell surfaces, where they regulate transport pathways of foreign molecules toward the cell. The structural and functional diversity of GAGs is largely attributed to varied sulfation patterns along the polymer chains, which makes understanding their molecular recognition mechanisms crucial. Molecular dynamics (MD) simulations, with their un-matched microscopic perspective, have the potential to be a reference tool for exploring the patterns responsible for biologically relevant interactions. However, the capability of molecular dynamics models (*i.e.*, force fields) used in biosimulations to accurately capture sulfation-specific interactions is not well established. In this work, we evaluate the performance of molecular dynamics force fields for sulfated GAGs by studying ion pairing of Ca^2+^ to sulfated moieties — N-methylsulfamate and methylsulfate — that resemble N- and O-sulfation found in GAGs, respectively. We tested nonpolarizable (CHARMM36 and GLYCAM06), explicitly polarizable (Drude and AMOEBA), and implicitly polarizable through charge scaling (prosECCo75 and GLYCAM-ECC75) force fields. The Ca–sulfamate/sulfate interaction free energy profiles obtained with the tested force fields were compared against reference ab initio molecular dynamics (AIMD) simulations. AIMD reveals that the preferential Ca^2+^ binding mode to sulfated GAG groups is solvent-shared pairing, and only the charge-scaled models agree satisfactorily with the AIMD data. All other force fields exhibit poorer performance, sometimes even qualitatively. Surprisingly, even explicitly polarizable force fields display a notable shortfall in their performance, attributed to difficulties in their optimization and possible inherent limitations in depicting high-charge-density ion interactions accurately. Finally, the underperforming force fields lead to unrealistic aggregation of sulfated saccharides, qualitatively distorting our understanding of the soft glycocalyx environment. Our results highlight the importance of accurately treating electronic polarization in MD simulations of sulfated GAGs and caution against over-reliance on currently available models without thorough validation and optimization.

## Introduction

Glycosaminoglycans (GAGs) are large linear polysaccharides, which exist either as free-floating molecules or as extended chains covalently bound to extracellular proteins known as proteoglycans, see Figure 1A. Along with glycoproteins, GAGs, and proteoglycans constitute the carbohydrate-rich cover layer of cell membranes of nearly all animal cells, referred to as the glycocalyx or extracellular matrix.^1,2^ The basic structural template of GAGs is relatively simple, consisting of repeating sequences of a uronic acid and an amino saccharide. ^1^ Despite such simplicity, the biological functions of GAGs are diverse and attributed to their structural heterogeneity, which stems from the saccharide type (*e.g.*, D-glucuronic acid vs. L-iduronic acid) and different amounts of sulfated groups, such as sulfates and sulfamates, unevenly distributed along the polysaccharide chains, ^3,4^ see Figure 1B. The unique arrangement of these sulfated groups, often termed the “sulfation code”, is responsible for numerous extracellular signaling events.^4^ GAGs were found to potentially interact with thousands of proteins,^5^ enabling and mediating a wide array of physiological processes.^6,7^ As a result, GAGs are involved in cell–cell communication and molecular recognition processes,^8–10^ and the development and progression of vascular problems,^11^ cancer,^12^ and Alzheimer’s disease.^13^

**Figure 1:**
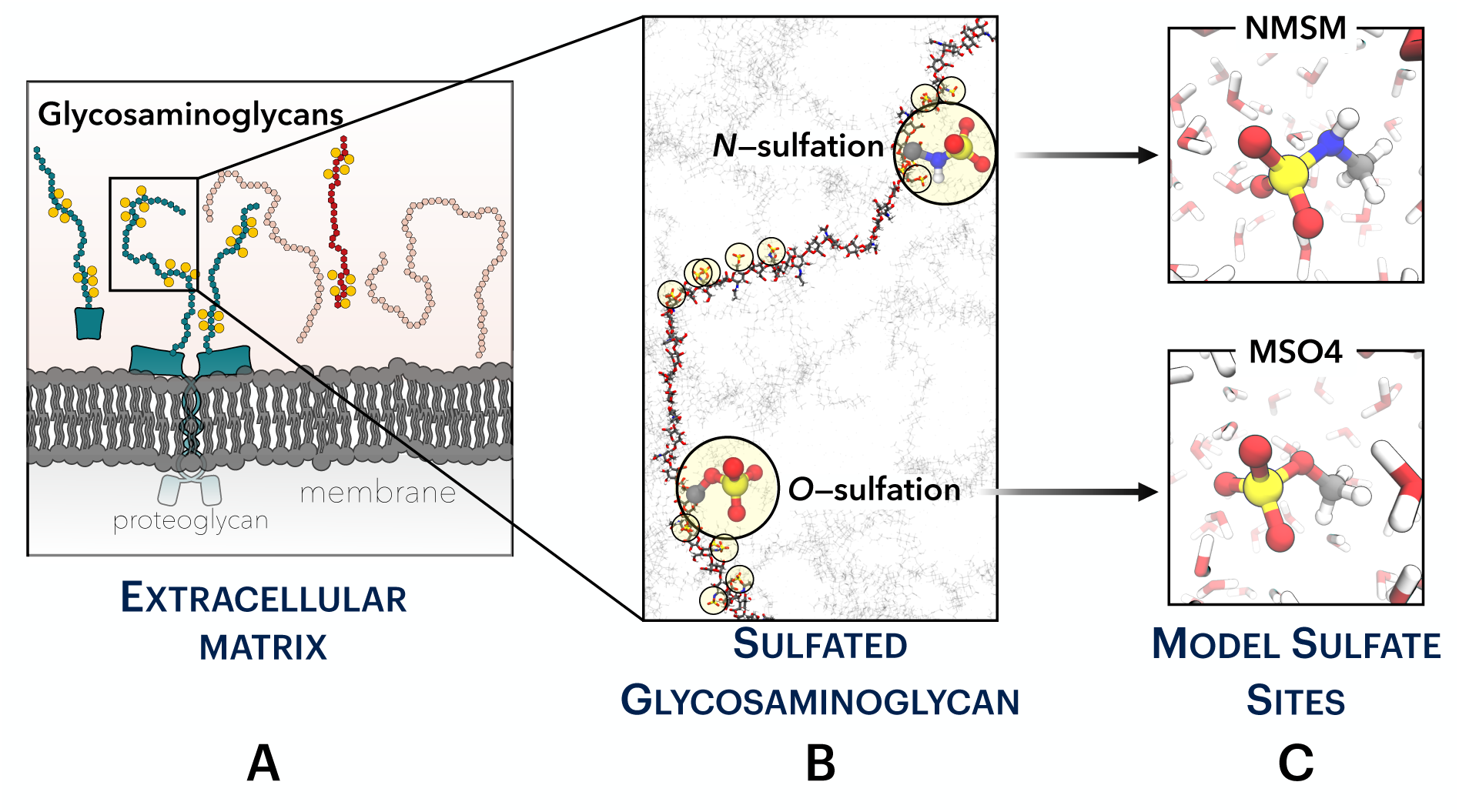
(A) Schematic illustration of the extracellular matrix emphasizing glycosaminoglycans (GAGs). GAGs exist in two primary forms: either as independent polysaccharide chains, e.g., hyaluronan (non-sulfated GAG; each monosaccharide is shown as a pink hexagon) and heparin (sulfated GAG; each monosaccharide is shown as a red hexagon), or as covalently attached segments of transmembrane or cleaved proteins, e.g., heparan sulfate and chondroitin sulfate (blue hexagons). All GAGs, except hyaluronan, feature numerous sulfate groups (yellow balls) along their structure. (B) Depiction of the sulfation types observed in GAG chains. (C) Model sulfate molecules — N-methylsulfamate (NMSM) and methylsulfate (MSO4) — resembling the N- and O-sulfation, respectively.

Studying the interaction of sulfated GAGs with other biomolecules is experimentally challenging due to the inherent variability in saccharide biological samples and the practi-cal difficulty of obtaining well-defined GAG sequences of relevant length.^14^ On top of that, traditional structural determination techniques typically struggle with studying interactions between GAGs and other moieties, such as proteins or ions. Molecular dynamics (MD) offers an excellent alternative that overcomes these experimental limitations. MD allows the inves-tigation of how sulfation defines the structure–activity relationship in a controlled molecular environment by imposing well-defined sulfation patterns. However, the application of MD to study GAGs remains relatively limited.^15^ The reason is that there are only a few ready-to-use sets of force field (FF) parameters for GAGs compatible with those for proteins, lipids, and other biomolecules in the extracellular space. Moreover, and perhaps most crucially, adequate experimental data for validating the accuracy of MD models of GAGs are scarce. This issue is especially pronounced for sulfated GAGs, whose experimental studies are costly and methodologically complex, particularly when short and unambiguously determined sequences are of interest.^16^ The lack of sufficient experimental validation raises questions about the accuracy of the available FFs, particularly atomistic ones, as they have not been thoroughly tested against experimental data, unlike their protein and lipid counterparts.

Nevertheless, the growing interest in studying GAGs is evident,^16^ particularly following the prominent role of glycans in the conformational dynamics of the receptor binding domain of SARS-CoV-2 spike protein.^17^ With computer simulations being at the forefront of glycan research,^18^ improving the accuracy of GAG models remains a priority goal. ^19^ Since all GAGs are negatively charged due to carboxyl and sulfate groups, benchmarking electrostatic interactions is a suitable approach to test the FFs’ performance. While the binding to carboxyl groups, including those of GAGs, has been addressed in several experimental and theoretical works,^20–23^ the binding to sulfated molecules has received less attention.^24^ GAGs possess N-sulfated and O-sulfated motifs, where a –SO_3_ group is covalently attached to either a nitrogen atom in an amino saccharide or an oxygen atom of a hydroxyl group, respectively. It has been proposed that the spatial conformation of GAGs is dictated by the distribution of these motifs and potentially their interactions with ions, ^25–27^ which are invariably present in biological environments.

Previous MD studies present the interactions of GAGs with Ca^2+^ as especially intriguing, given their presumed role in regulating GAG–protein binding.^28^ This regulation was attributed to the capacity of divalent cations for multidentate or “bridging” enabling interactions with multiple negatively charged groups simultaneously. ^29^ Such interactions, along with fundamental electrostatic screening effects, were presented as determining the flexibility and functional state of GAGs within cellular environments.^30,31^ However, Ca^2+^ simulations are prone to artifacts if electrostatic interactions are not appropriately handled. ^32–34^ This makes it unclear whether previous MD results report the real behavior of GAGs in the presence of Ca^2+^ or whether they are caused by artifacts related to ill-defined electrostatic interactions.^33^

In this study, we address the aforementioned scientific and methodological challenges associated with GAG simulations. We advocate for the use of ab initio molecular dynamics (AIMD) simulations as reference data to benchmark and improve molecular dynamics FFs for GAGs. AIMD has previously been proven as a robust method to verify the accuracy of molecular dynamics models for monoatomic ions and small molecules. ^20,35^ We utilize AIMD simulations in combination with enhanced sampling to assess the binding free energy of Ca^2+^ to small sulfated molecules resembling N- and O-sulfated saccharides. Then, we evaluate the performance of existing molecular dynamics FFs against AIMD data, aiming to discern how they capture intricate GAG–cation interactions. We examine all-atom non-polarizable FFs (CHARMM36^36^ and GLYCAM06^37^), all-atom implicitly polarizable FFs through the scaling of partial charges (prosECCo75^38^ and GLYCAM-ECC75 — derivatives of CHARMM36 and GLYCAM06, respectively), and all-atom explicitly polarizable FFs (Drude^39^ and AMOEBA^40^). Hereafter, we refer to these as force field molecular dynamics (FFMD) simulations. Our findings reveal significant disparities in the existing FFs’ abilities to accurately depict ionic pairing, underscoring the need to use charge-scaled FFs and to further develop or refine existing models for GAGs. This refinement is essential for ensuring the accuracy of future large-scale simulations involving GAGs, as well as other sulfated molecular moieties (e.g., proteins, lipids, and drugs) that participate in biological processes.

## Methods

In this work, we performed extensive free energy simulations of Ca^2+^–sulfamate/sulfate interactions using several molecular moieties and corresponding simulation models. We also modeled sulfated GAG disaccharides to study their behavior in an aqueous solution with calcium cations via unbiased MD simulations.

### Simulation Systems to Study Sulfation in Glycosaminoglycans

We used simplified systems as proxies to investigate sulfation in GAGs, and DFT-based AIMD served as a reference for benchmarking sulfate and sulfamate parameters in FFMD models. The negatively charged N-methylsulfamate (NMSM — CH_3_NHSO_3_*^−^*) and methyl-sulfate (MSO4 — CH_3_OSO_3_*^−^*) molecular ions were chosen to model the N- and O-sulfation of GAGs, respectively, see Figure 1C. NMSM contains a sulfamate –N–SO_3_*^−^* group, while MSO4 features a sulfate –O–SO_3_*^−^*group, both with a methyl residue. Modeling these molecules was a necessary choice since performing extensive free energy simulations on sulfated monosaccharides is currently unfeasible at the ab initio level. Nevertheless, we modeled single-sulfated (either N-sulfated or 6-O-sulfated) N-acetyl-d-glucosamine at the FF level to test the performance of our small molecules in resembling sulfation in GAGs. Finally, we also performed FFMD simulations with sulfated disaccharides composed of L-iduronic acid (IdoA) and N-acetyl-d-glucosamine (GlcNAc) (*α*-IdoA-*α*(1,4)-GlcNAc), a key fragment in sulfated region of heparan sulfate,^4^ with either N- or O6-site sulfation of the amino unit.

The procedure of building the molecules and obtaining their topologies is described in section S1 in the Supporting Information (SI). All the topologies, input configurations, and additional files necessary for reproducing our data are openly available in Zenodo at DOI: 10.5281/zenodo.10036627

### Ab Initio Molecular Dynamics Simulations with NMSM and MSO4 Molecules

The free energy of calcium binding to NMSM and MSO4 anions was determined using the DFT-AIMD approach performed by the Quickstep module of CP2K 7.1^41,42^ software. The AIMD system contained a single Ca^2+^, one molecular anion (MSO4 or NMSM), and 128 water molecules. The resulting system net +1 charge was compensated by a uniform background charge of the opposite sign. The systems were propagated with 0.5 fs time step in the *NVT* ensemble at 300 K using three-dimensional periodic boundary conditions. Every initial system configuration underwent equilibration under the Langevin thermostat^43^ with a time constant of 50 fs. Subsequently, the production runs employed the stochastic velocity rescaling thermostat^44^ with a time constant of 1 ps.

The electronic structure was calculated on-the-fly using the revPBE-D3 GGA density functional.^45–48^ To avoid spurious overpolarization of monoatomic cations, the D3 correction was turned off for all pairs containing Ca^2+^.^49^ For comparison, the results with the potentially problematic^49^ D3 correction enabled for all atomic pairs are provided in section S3.1 in the SI. The valence Kohn–Sham orbitals were expanded into the TZV2P basis set for H, N, O, C, S, and TZV2P-MOLOPT-SR-GTH for Ca,^50^ while the core orbitals were described using the Goedecker–Teter–Hutter (GTH) pseudopotentials.^51^ The self-consistent field convergence criterion was set to 10*^−^*^5^ Ha. Additionally, an auxiliary plane-wave basis with a 400 Ry energy cutoff was used to represent the electronic density within the GPW scheme.^52^

### Force Field Molecular Dynamics Simulations with NMSM and MSO4 Molecules

Using NMSM and MSO4 molecules, we evaluated several FFs for their ability to accurately capture the free energy of Ca^2+^–sulfamate/sulfate interactions. These FFs fall into three categories: nonpolarizable — CHARMM36 (including its current default version with NBFIX (i.e., Non-Bonded Fix) corrections) and GLYCAM06 (version j), implicitly polarizable via charge scaling — prosECCo75 and GLYCAM-ECC75, and explicitly polarizable — Drude and AMOEBA. We used typical force-field-specific molecular dynamics protocols for all of these models, see Sections S1 and S2 in the SI for the key FF parameters, simulation details, and software used.^53–59^ Identically to AIMD, our systems contained a single Ca^2+^, one molecular anion (MSO4 or NMSM), and 128 water molecules, and the system net charge was compensated by a uniform background charge. The box size, unless stated otherwise, was set to around 1.59 nm each side, which restricts the cutoffs for electrostatic and Lennard–Jones interactions to 0.7 nm, which is smaller than common 1–1.2 nm, see Section S2 for details. Additionally, we performed FFMD simulations using a larger cubic box, with *∼*31.3 Å on each side and containing 1024 water molecules. In this case, we used force-field recommended cutoffs, and we were able to estimate the absolute binding free energy of calcium to the studied anions. Using these larger systems, we also discarded potential artifacts due to the small cutoffs imposed in smaller systems, see Section S3.5. Finally, we performed free energy simulations where both NMSM and MSO4 molecules were simultaneously present with a Ca^2+^ cation and 2048 water molecules.

While we adopted already existing parameters for simulations with nonpolarizable and explicitly polarizable FFs, this study derived the prosECCo75 and GLYCAM-ECC75 pa-rameters for sulfated saccharides and corresponding model molecules. The idea behind pros-ECCo75 and GLYCAM-ECC75 is to incorporate electronic polarization in a mean-field way by scaling down atomic partial charges.^33^ This approach — known as Molecular Dynamics in Electronic Continuum, Electronic Continuum Theory, or Electronic Continuum Correction (ECC)^60–62^ — suggests that the scaling factor should be equal to the reciprocal square root of the high-frequency dielectric constant of the surrounding medium, *i.e.*, 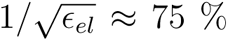 in the case of aqueous solutions and most biological environments. ^33^ Very recent extensive AIMD simulations confirm that the scaling factor indeed lies in the 75 % to 80 % range.^63^ However, the optimal value still remains debated since it is influenced by the dielectric constant of the chosen water model,^64^ partially included electronic polarization in the original parameterizations,^33^ and the specific properties of the system under study.^65^

Nevertheless, there are numerous examples where the ECC approach with the theoretically sound 75 % scaling improved the quality of FFMD. These examples include simulations of monoatomic ions, ^20,32,66^ small ligands,^22,67^ biomolecules,^23^ lipid membranes,^68^ and even metal-oxide surfaces.^69^ We have recently developed the prosECCo75 model, a 75 %-scaling ECC patch to the CHARMM36 FF.^38^ Our prosECCo75 patch replaces in a physically justified manner the existing NBFIX correction that adds repulsive terms in the Lennard–Jones to prevent the excessive solute–solute association in CHARMM36 simulations.^70^ However, the current iteration of the prosECCo75 model does not include ECC charges for sulfation. Thus, our work fills this gap by introducing prosECCo75-compatible parameters for sulfamate and sulfate groups, Table 1.

**Table 1:**
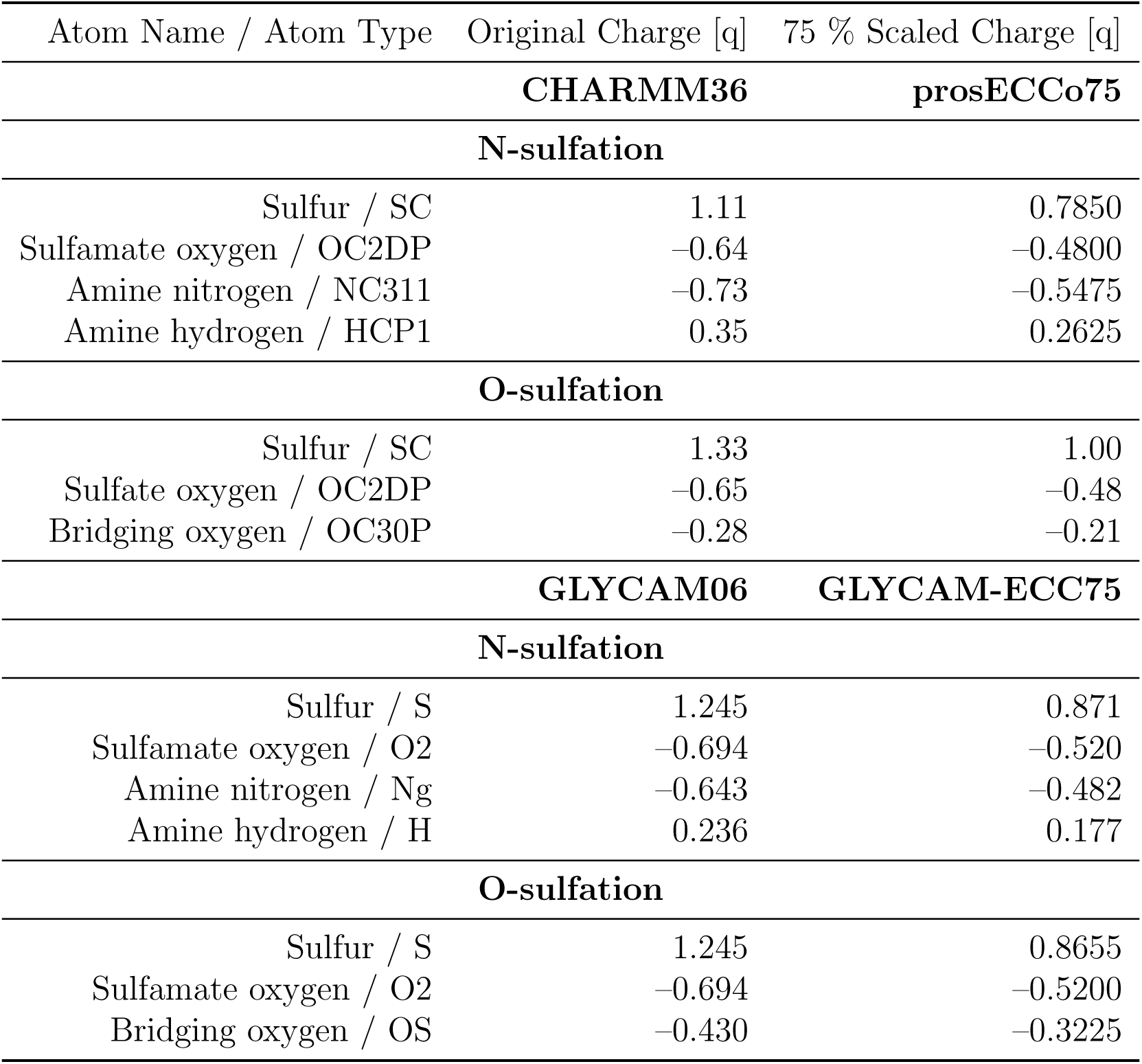
Partial atomic charges and atomic types in sulfate and sulfamate groups of GAGs for CHARMM36 and GLYCAM06 force fields and their charge-scaled derivatives prosECCo75 and GLYCAM-ECC75, respectively.

Similarly, ECC subversions of AMBER and AMBER-compatible FFs have been previously tested.^71,72^ To the best of our knowledge, however, there have been no attempts to model sulfated saccharides by applying charge scaling to the GLYCAM06 FF. Therefore, we have adopted the ECC approach for the GLYCAM model, resulting in a variant termed GLYCAM-ECC75, see again Table 1. Note that the charge scaling is applied to only ionic groups,^33,38^ which are typically as distant as possible from dihedrals critical to structural conformations, such as saccharide ring puckering. ^38^ This strategy is designed to minimize alterations to the well-established CHARMM36 and GLYCAM06 models.

### Force Field Molecular Dynamics Simulations of Saccharide Solutions

To validate the use of NMSM and MSO4 molecules as mimics to study sulfation in GAGs, we computed FFMD free energy profiles for monosaccharides. The systems contained either one N-sulfated N-acetyl-d-glucosamine or one 6-O-sulfated N-acetyl-d-glucosamine, together with one Ca^2+^ and 1024 water molecules.

We also modeled disaccharides composed of L-iduronic acid (IdoA) and N-acetyl-d-glucosamine (GlcNAc) (*α*-IdoA-*α*(1,4)-GlcNAc) sulfated in either N- or O6-site of GlcNAc. We used them for unbiased simulated systems consisting of 16 disaccharides (N- or O-sulfated) and 16 Ca^2+^ cations solvated in a cubic box with *≈* 4 nm each side and containing *≈* 1850 water molecules. Given that each disaccharide features one carboxyl group in the IdoA and one sulfate/sulfamate group in the GlcNAc, the overall charge of the system was zero. We only tested nonpolarizable and implicitly polarizable FFs. These disaccharide systems were shortly minimized using the steepest descent algorithm and then simulated for 200 ns in the *NPT* ensemble, which was sufficient for the converged results. The first 100 ns served as the equilibration period, and the last 100 ns were used for the analysis,

Initial coordinates and topologies of saccharide structures for CHARMM36 simulations were obtained using the Glycan Modeler within CHARMM-GUI utility.^73,74^ GAG builder at GLYCAM-Web (https://glycam.org/gag/)^75^ was employed to generate parameters for GLYCAM06. The GLYCAM topologies were then translated from AMBER to GROMACS format using the ACPYPE tool.^76^ Further simulation details are given in Section S2.

### Umbrella Sampling Simulations

Umbrella sampling simulations, followed by potential of mean force (PMF) calculations, were performed to estimate the interaction free energies between Ca^2+^ and the studied molecules, *i.e.*, NMSM, MSO4, and monosaccharides, for the above FFMD and AIMD systems. Unless stated otherwise, the procedure for generating umbrella sampling configurations, performing corresponding simulations, and post-processing trajectories was always the same. We chose the Ca^2+^–sulfur distance as the collective variable since the sulfur atom provides a suitable and consistent reference for both NMSM and MSO4 molecules and the corresponding monosaccharides, see Section S3.3. From a FFMD pulling simulation, we generated initial configurations spanning various Ca^2+^–sulfur distances. The reference distances and corresponding harmonic force constants are summarized in Section S2.5 in the SI. These configurations were then equilibrated for 50 ns each employing the prosECCo75 model. The equilibrated structures served as the initial configurations for all subsequent AIMD and FFMD free energy simulations.

All umbrella sampling simulations were performed in the *NVT* ensemble in a cubic box. Although we observed minor deviations in the resulting box size from *NPT* simulations with different FFs, these deviations have no impact on the free energy profiles, see Section S3.4 in the SI. Therefore, the reported results use an identical box size for all umbrella sampling configurations for a given molecule (NMSM, MSO4, or monosaccharide) and system composition. The used box size corresponded to the average box size in a *NPT* simulation of the corresponding system using the prosECCo75 FF. In AIMD, each umbrella sampling window was equilibrated for 5 ps, followed by a 200 ps production run. In FFMD simulations, except those with AMOEBA model, each umbrella window simulation was 50 ns long, with the first 10 ns serving as an equilibration period. In AMOEBA simulations, each umbrella sampling window was simulated for 11 ns, discounting the initial 1 ns as equilibration time.

The corresponding PMF profiles were extracted using gmx wham utility implemented in GROMACS.^77^ To estimate errors, we employed bootstrap analysis using 100 bootstrap samples as implemented in the same utility.^77^ Our graphical representations of the free energy profiles omit the error bars from FFMD simulations when they are smaller than the thickness of the plotted free energy lines. We applied the volume entropy correction to all PMF profiles^78^ (+2*k_B_T* ln(*r*)). Then, in the case of systems with 128 water molecules, we shifted the PMF profiles to zero at the energy minimum that corresponds to the solvent-shared ion pairing. This alignment allows direct comparison of free energy differences between the contact ion and solvent-shared pairings across both AIMD and FFMD simulations. In order to estimate the absolute binding free energies in the systems with 1024 water molecules, the energy profiles were shifted to zero at the largest distances, where Ca^2+^ does not interact with a NMSM/MSO4/monosaccharide.

Finally, we also performed multiple checks on selected systems to validate our umbrella sampling methodology and corresponding results, by checking the effect of the box size (Section S3.4), missing counterion (Section S3.6), imposed cutoff and treatment of electrostatic and Lennard–Jones interactions (Sections S3.7 and S3.8), and water model (Sections S3.9 on the free energy profiles. All of these checks confirmed that the main conclusions of this work are not affected by any of the aforementioned factors.

### Accelerated Weight Histogram Method

We used the Accelerated Weight Histogram (AWH) method implemented in GROMACS^79^ to calculate two-dimensional free energy profiles of Ca^2+^ interactions with both NMSM and MSO4 simultaneously present. In this case, we tested only CHARMM36-NBFIX and prosECCo75 FFs. The simulation setup comprised one NMSM, one MSO4 molecule, a single Ca^2+^, and 2048 water molecules. Simulations were performed in the *NPT* ensemble, using the recommended values for all simulation parameters, including cutoffs, as prescribed by the FFs tested. We used two reaction coordinates — Ca^2+^–sulfur distances to both NMSM and MSO4 — over a range from 2.7 to 15 Å. Each AWH simulation was 1 *µ*s long. The two-dimensional volume entropy correction was applied by summing independently the two one-dimensional corrections.

We also used AWH to verify the convergence of our umbrella sampling simulations and to ensure that the applied force constants do not stifle conformational sampling. We calculated the free energy profiles for selected systems with 128 water molecules. These simulations were conducted along the same reaction coordinate (Ca^2+^–sulfur distance) spanning a range from 2.7 to 7 Å. Each AWH simulation was 50 ns long. The obtained AWH profiles were found indistinguishable from those acquired using the umbrella sampling simulations, see Section S3.10 in the SI.

For all AWH calculations, the initial target distribution was chosen to be uniform. The initial diffusion constant and error estimate were set to 5 *·* 10*^−^*^5^ nm^2^ *·* ps and 5 kJ *·* mol*^−^*^1^, respectively. The force constant of the umbrella potential was set to 100, 000 kJ *·* mol*^−^*^1^ *·* nm*^−^*^2^. The comprehensive technical details of the AWH method can be found in the original article.^80^

## Results and Discussion

### Small Sulfated Molecules Accurately Resemble Sulfation in Glycosaminoglycans

We performed free energy FFMD simulations on sulfated monosaccharides and compared them with those of NMSM and MSO4 to validate the use of the latter as simplified models of sulfation in GAGs. Our simulations show that the obtained R–N/O–SO_3_*^−^*–Ca^2+^ (R = CH_3_ or saccharide) binding free energy profiles are virtually identical, see section S3.2 in the Supporting Information (SI). Therefore, NMSM and MSO4 molecules are well-suited for the purposes of this work.

### Calcium Preferentially Interacts with NMSM and MSO4 via Solvent-Shared Pairing

Figure 2 (top) shows the Ca^2+^–NMSM (left) and Ca^2+^–MSO4 (right) interaction free energies obtained from the AIMD simulations. The similarity in the energy profiles is remarkable, suggesting that the influence of the bridging atom (nitrogen in NMSM and oxygen in MSO4) on Ca^2+^ coordination can be neglected. Therefore, further discussion of the results will broadly cover both N- and O-sulfation.

**Figure 2:**
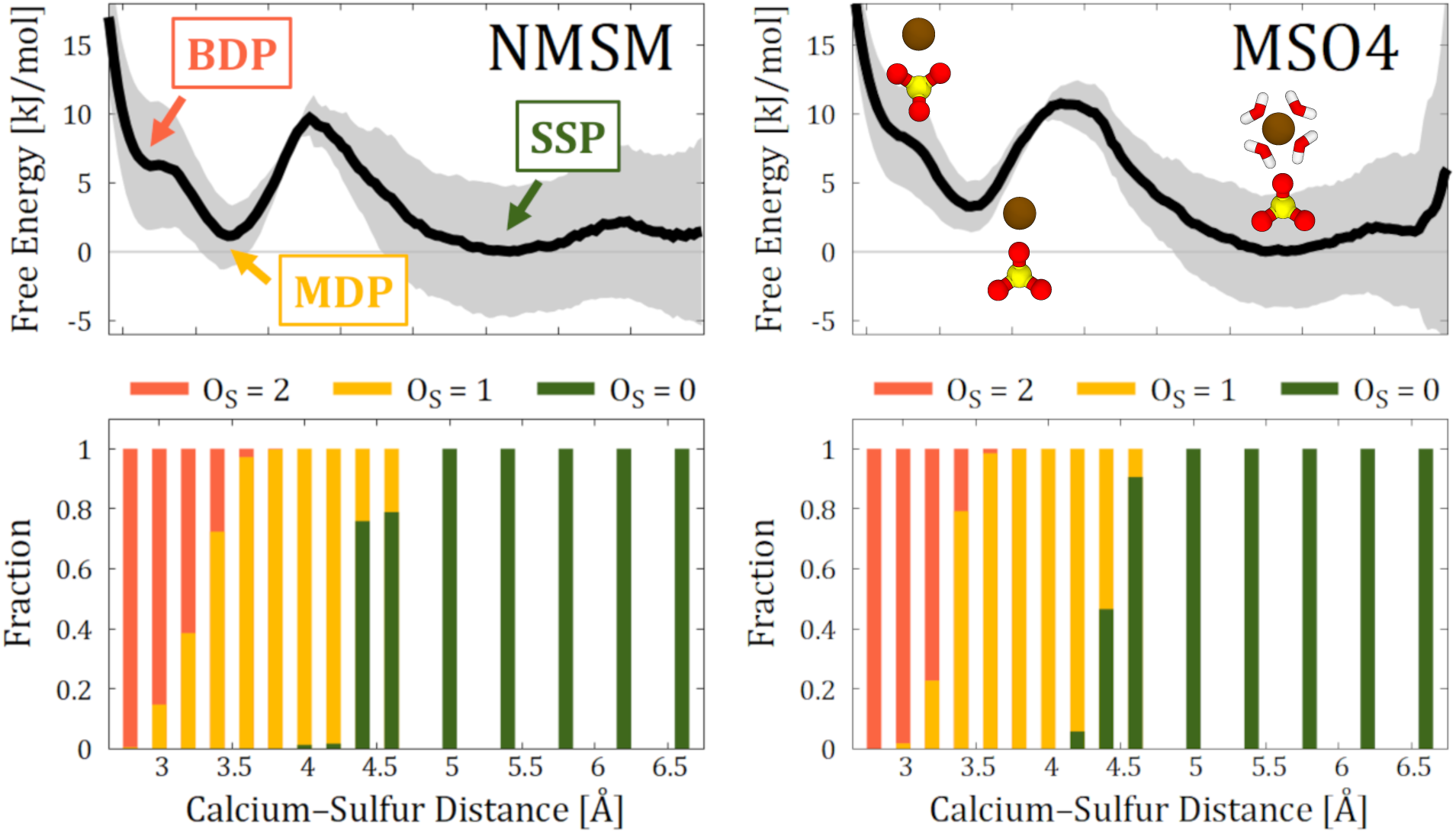
(top) Free energy profiles as a function of the Ca^2+^–S distance, derived from AIMD umbrella sampling simulations. Three distinct binding modes are identified: bidentate pairing (BDP), monodentate pairing (MDP), and solvent-shared pairing (SSP). (bottom) Fractions of sulfamate/sulfate oxygens (O_S_) within the first solvation shell of Ca^2+^ along the distances defined by the umbrella sampling windows. The first solvation shell is defined by the first minimum in the Ca^2+^–water radial distribution function and located at 3.35 Å.

Three distinct regions can be found in the free energy profiles. The first noticeable feature is located approximately at around 3.2 Å, which is, at best, a higher-energy metastable state that corresponds to the bidentate ion pairing (BDP) of the Ca^2+^, which coordinates with two oxygens of the sulfamate/sulfate group. Then we observe a local minimum at around 3.7 Å corresponding to the monodentate ion pairing (MDP), *i.e.*, Ca^2+^ is coordinated mostly to only one of the oxygens. Finally, the region at *≈* 5.0–6.0 Å denotes solvent-shared ion pairing (SSP), characterized by the intact hydration shells of interacting ions. The structural characteristics of these binding modes become evident when examining the number of sulfamate/sulfate oxygen atoms in the first solvation shell of Ca^2+^, see Figure 2 (bottom). Essentially, the hydration water oxygens are substituted by those from the sulfamate/sulfate group in monodentate and bidentate binding modes, and the total oxygen count remains around *≈* 6.3–6.8.

Our findings clearly demonstrate that the dehydration of Ca^2+^ does not fully determine its interaction with sulfamate/sulfate groups, as the strongest pairing happens with intact solvation shells in both cation and anion. This contrasts with the monodentate ion pairing preferred significantly by other anions, such as those with carboxyl groups,^20,21,26^ which are also present in saccharides. Despite previous suggestions of solvent-shared Ca^2+^ binding to GAGs,^26^ there is no consensus in the literature on the predominance of this interaction mode in binding to sulfamate/sulfate groups. ^28,29,81^ Our AIMD simulations suggest that both monodentate and solvent-shared pairing of Ca^2+^ to sulfamate/sulfate groups are plausible at biologically relevant conditions. The solvent-shared mode is slightly favored over the monodentate mode by approximately 1–3.5 kJ/mol, while the bidentate mode is much less likely, being higher in energy by around 6–9 kJ/mol when compared to solvent-shared pairing. The free energy barrier between the solvent-shared and monodentate states is *≈* 10 kJ/mol.

It is fair to acknowledge non-negligible errors in our AIMD energy estimates. Also, the energy increase at the furthest distances for MSO4 likely results from fewer sampling due to a single umbrella sampling window covering this region. Nevertheless, the overall high similarity between the two profiles lends credibility to our analysis and the subsequent conclusions drawn from the data.

### Common Nonpolarizable Force Fields Overestimate Cation–Sulfamate and Cation–Sulfate Interactions

CHARMM36^36,82–85^ and GLYCAM06^37,86^ are currently the most popular FFs for saccharide simulations, including GAGs, owing that to their satisfactory performance and computational efficiency compared to polarizable FFMD or AIMD. The GLYCAM06 FF is designed to be compatible with the AMBER family of models, ^87,88^ while CHARMM36 saccharides are compatible with all other biomolecules provided by the CHARMM family, including proteins, lipids, DNA, and other biomolecules. Our data shows that both CHARMM36 and GLYCAM06 tend to overestimate the strength of contact ion pairing, heavily favoring the monodentate binding, see Figure 3. This issue stems from the overestimated electrostatic interactions typical for nonpolarizable FFs.^23,24,33,62^

**Figure 3:**
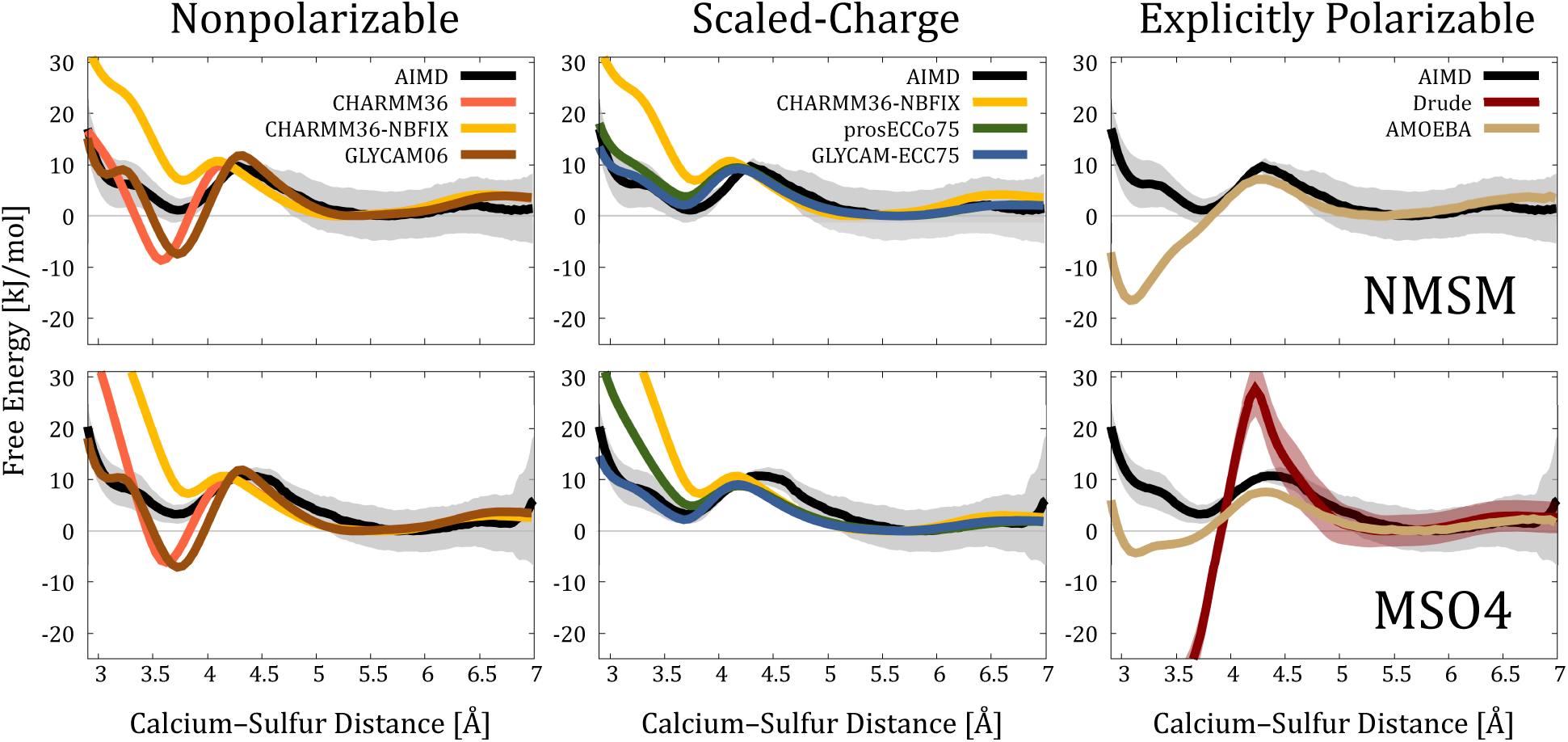
Comparison of the interaction free energy profiles obtained from AIMD and FFMD simulations. The FFMD profiles are grouped according to the treatment of electronic polarization. CHARMM36-NBFIX profiles are shown twice, as they are added to the “Scaled-Charge force fields” block due to the similar reasoning in NBFIX and scaled-charge approaches. The Drude profile for Ca^2+^–NMSM binding is not given because of the numerical instability in the corresponding simulations caused by the overpolarization. An alternative version of this figure with adjusted vertical scales for better per-force-field visualization is given in the SI, Section S3.11.

To address the spurious electrostatic interactions, CHARMM36 model incorporates the so-called NBFIX correction, which modifies pair-specific Lennard–Jones parameters to mitigate excessive electrostatic interactions.^70^ This correction is a default feature in the latest CHARMM36m FF,^36^ and it also contains tailored parameters to fine-tune Ca^2+^ interactions with sulfamate and sulfate groups. Therefore, we performed additional simulations with NBFIX correction turned on, referred to as CHARMM36-NBFIX (yellow curves in the left and central panels of the Figure 3). The results from these CHARMM36-NBFIX simulations suggest that the introduced van der Waals repulsion is overly strong, now leading to the underrepresentation of monodentate binding and significantly hindered bidentate binding.

The inherent issue with nonpolarizable parameterizations emerges already in NMSM/MSO4– water interactions. We calculated the radial distribution functions (RDFs) of water hydrogens (H_w_) around the sulfamate/sulfonate oxygens (O_S_) from the umbrella sampling simulation window where the calcium cation and the studied molecules are at their maximum distance, basically excluding the contribution of bound Ca^2+^, see Figure 4. For nonpolarizable FFs, our results indicate a pronounced overstructuring of water molecules around the sulfamate/sulfate groups compared to the AIMD data. A similar overstructuring problem was also observed for Ca^2+^.^20,32^ Therefore, merely adjusting dispersion cross-interactions, as in NBFIX, is insufficient to fix inadequate electrostatic screening as it does not directly impact NMSM/MSO4–water or Ca^2+^–water interactions.

**Figure 4:**
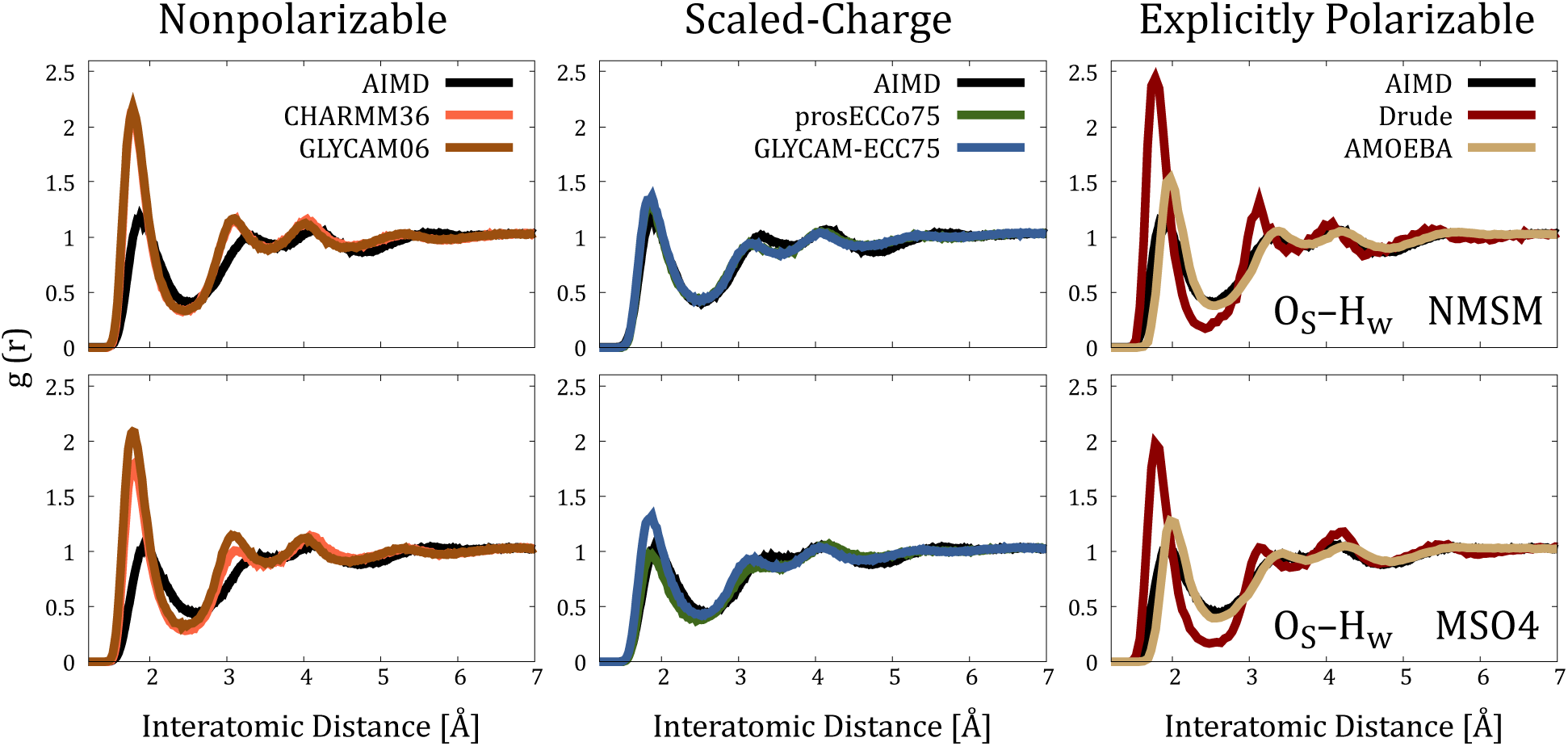
Comparison of O_S_–H_w_ radial distribution functions (RDFs) collected from AIMD and FFMD simulations. Additional RDFs are summarized in Section S3.12 in the SI.

### Scaling Charges Provides Prominent Results

Scaling partial atomic charges offers a robust mean-field approach to account for the missing electronic polarization without the need for more computationally demanding and harder-to-parameterize polarizable FFs.^60–62^ Both scaled-charge derivatives considered, one from CHARMM36 and the other from GLYCAM06 — prosECCo75 and GLYCAM-ECC75, re-spectively — provide much closer agreement with the AIMD data, see Figure 3. They greatly capture the energy differences between monodentate and solvent-shared modes, as well as the height of the energy barrier, when compared to AIMD. The bidentate pairing is better reproduced by GLYCAM-ECC75, particularly in the case of Ca^2+^–MSO4 pairing.

Pursuing our analysis for nonpolarizable FFs, we again calculated the sulfamate/sulfate– water RDFs. Both prosECCo75 and GLYCAM-ECC75 models demonstrate an excellent agreement with the AIMD data, effectively capturing all features of anion hydration. It is worth noting that this level of agreement is achieved solely by scaling the partial charges of ionic groups without modifying the Lennard–Jones potentials, c.f. Table 1. Additionally, incorporating electronic polarization substantially improves the representation of the hydration shell of calcium,^21,34^ as we show in Section S3.12 in the SI, altogether improving the description of NMSM/MSO4–calcium interactions.

### Explicitly Polarizable Force Fields Significantly Underperform

FFs incorporating electronic polarization explicitly are often considered superior to the non-polarizable models. There are currently two popular choices of such FFs for biosimulations: Drude^39^ and AMOEBA,^40^ each incorporating electronic polarization in a different way. The Drude FF uses negatively charged auxiliary particles attached to non-hydrogen atoms by a harmonic spring, which mimic the charge fluctuations under the influence of an electric field. In AMOEBA FF, the electronic response is achieved by self-consistently calculating classical dipole moments due to multipoles (up to quadrupole) and isotropic atomic polarizabilities assigned to each polarizable atom.

Despite their potential, the Drude and AMOEBA FFs have not been widely adopted in GAG simulations. The main obstacles are the lack of well-established parameters, although generic Drude parameters have been proposed, ^89–91^ and higher computational costs, which can be critical when simulating GAGs at biologically relevant lengths and time scales. Nevertheless, the community eagerly awaits the application and development of explicitly polarizable FFs for GAGs, ^18^ especially the sulfated ones, given their anticipated potentially improved accuracy in describing the charged nature of GAGs.

Contrary to initial expectations, Drude and AMOEBA perform worse than traditional nonpolarizable FFs in simulating Ca^2+^ binding to NMSM and MSO4 molecules, Figure 3. Both models significantly overestimate the propensity for bidentate pairing. AMOEBA favors the bidentate binding over solvent-shared by *≈* 17 kJ/mol for NMSM and about 4 kJ/mol for MSO4. Drude suggests a clearly excessively stronger preference for the bidentate mode in MSO4 binding (almost 70 kJ/mol, see Figure S19), while Ca^2+^ interactions with NMSM are so strong that they compromise the geometry of the NMSM molecule, see Section S2.3 in the SI.

Such a subpar performance is likely rooted in the lack of specific parameter tuning for moieties with a high charge density such as Ca^2+^ and oxyanions.^22,92–94^ Several studies have pointed out the overpolarization of Ca^2+^ in Drude simulations and the necessity for additional terms like NBFIX or NBTHOLE.^92–94^ For AMOEBA, an improvement might come with future developments within AMOEBA+,^95^ which incorporates short-range charge penetration and charge transfer terms into the model. AMOEBA+ parameters for sulfamate/sulfate groups are not available at the moment.

Further elaborating on challenges with explicitly polarizable FFs, we revisited the corresponding anion–water RDFs. As illustrated in Figure 4, Drude predicts excessively rigid hydration shells around sulfamate/sulfate groups, similarly to nonpolarizable FFs. In contrast, AMOEBA provides a reasonable level of accuracy, albeit with the first peak being noticeably displaced to shorter distances. Thus, the overestimation of Ca^2+^ binding in AMOEBA could likely be improved/rectified by tuning Ca^2+^–sulfamate/sulfate interactions. Note that the first hydration shell around NMSM is tighter than around MSO4 in AMOEBA simulations, because the NMSM parameterization is more polar, potentially leading to stronger Ca^2+^ binding.

### Solvent-Shared Calcium Can Mediate Intra- and Intermolecular Interactions

We investigated whether Ca^2+^ can mediate interactions between two sulfamate/sulfate groups. We utilized CHARMM36-NBFIX and prosECCo75 FFs to generate two-dimensional free energy profiles using the Accelerated Weight Histogram (AWH) method. Our results unambiguously demonstrate that in simulations employing either FF, the combined solvent-shared mode emerges at the symmetric 5–6 Å distances as the most energetically favorable configuration, see Figure 5. This results in an energy minima of approximately –8 kJ/mol and –4.5 kJ/mol for CHARMM36-NBFIX and prosECCo75, respectively. Consequently, our findings indicate that Ca^2+^ can foster interactions between GAGs or interactions with other biomolecules possessing negatively charged groups.

**Figure 5:**
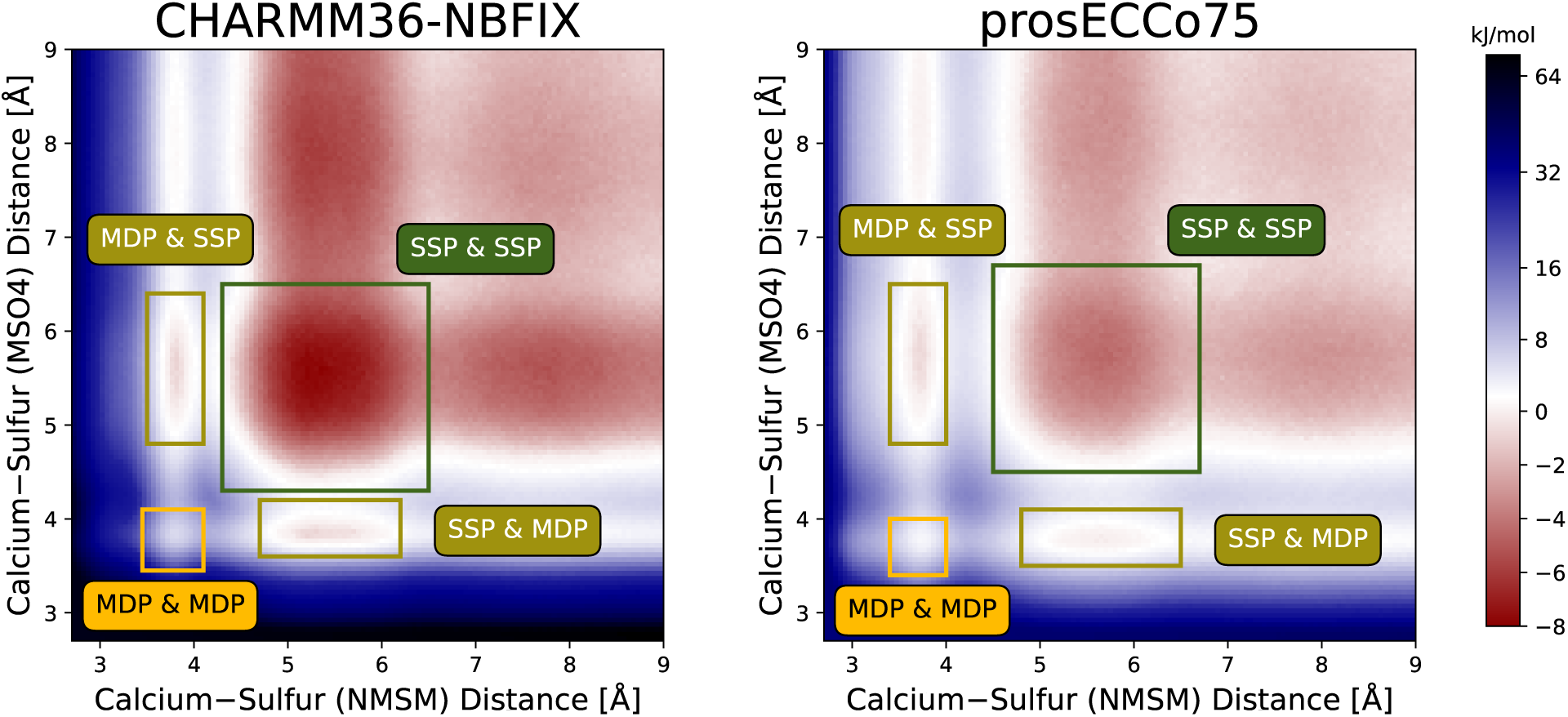
Two-dimensional free energy profiles of Ca^2+^ concurrent binding to NMSM and MSO4 molecules. The energy profiles were obtained from AWH simulations with CHARMM36-NBFIX and pros-ECCo75 force fields. The combined binding modes to both NMSM and MSO4 molecules are highlighted by colored rectangles. The error estimated by gmx awh tool is 0.32 kJ/mol.

### prosECCo75 and CHARMM-NBFIX Prevent Unphysical Aggregation of Calcium–Saccharide Solutions

We examined how the (in)accurate description of Ca^2+^–sulfamate/sulfate interactions influence the structure of saccharide solutions. For this purpose, we selected a simple and representative example of disaccharides (sulfated either at N- or O-position) solvated in water with Ca^2+^. We tested only nonpolarizable and scaled-charge FFs, as explicitly polarizable models have demonstrated inadequate performance already for NMSM and MSO4. Our analysis focused on monitoring the formation of molecular clusters, typically driven by strong contact ion pairing and inadequate saccharide–saccharide interactions. ^96^ Such phenomena are frequently observed when there is an imbalance between solute–solute and solute–solvent interactions, resulting in the unphysical aggregation of experimentally highly soluble species.

Indeed, simulations with CHARMM36 and GLYCAM06 FFs, both showing strong monodentate pairing between Ca^2+^ and NMSM/MSO4, revealed significant clustering, see Figure 6A. The issue was especially pronounced with the GLYCAM06 model, where the system predominantly formed a single large cluster. Interestingly, the charge scaling did not always fully solve this issue. Despite GLYCAM-ECC75 showing excellent results in reproducing AIMD Ca^2+^–sulfamate/sulfate binding free energies, even better than prosECCo75, especially in the bidentate binding region, simulations using GLYCAM-ECC75 model still showed a noticeable cluster formation not present in prosECCo75, see Figure 6B. Earlier studies have already noted the issue of excessive aggregation in GLYCAM06 simulations;^96,97^ however, a consistent solution to this problem has yet to be established. While incorporating the ECC framework improves electrostatic interactions with sulfamate/sulfonate groups in saccharides, c.f. Figure 3, imbalances in other interactions, such as those between saccharide molecules themselves, may still persist. This highlights the importance of the subtle balance between saccharide–saccharide, saccharide–cation, saccharide–water, cation–water, and water–water interactions, all contributing to the overall not always intuitive behavior.

**Figure 6:**
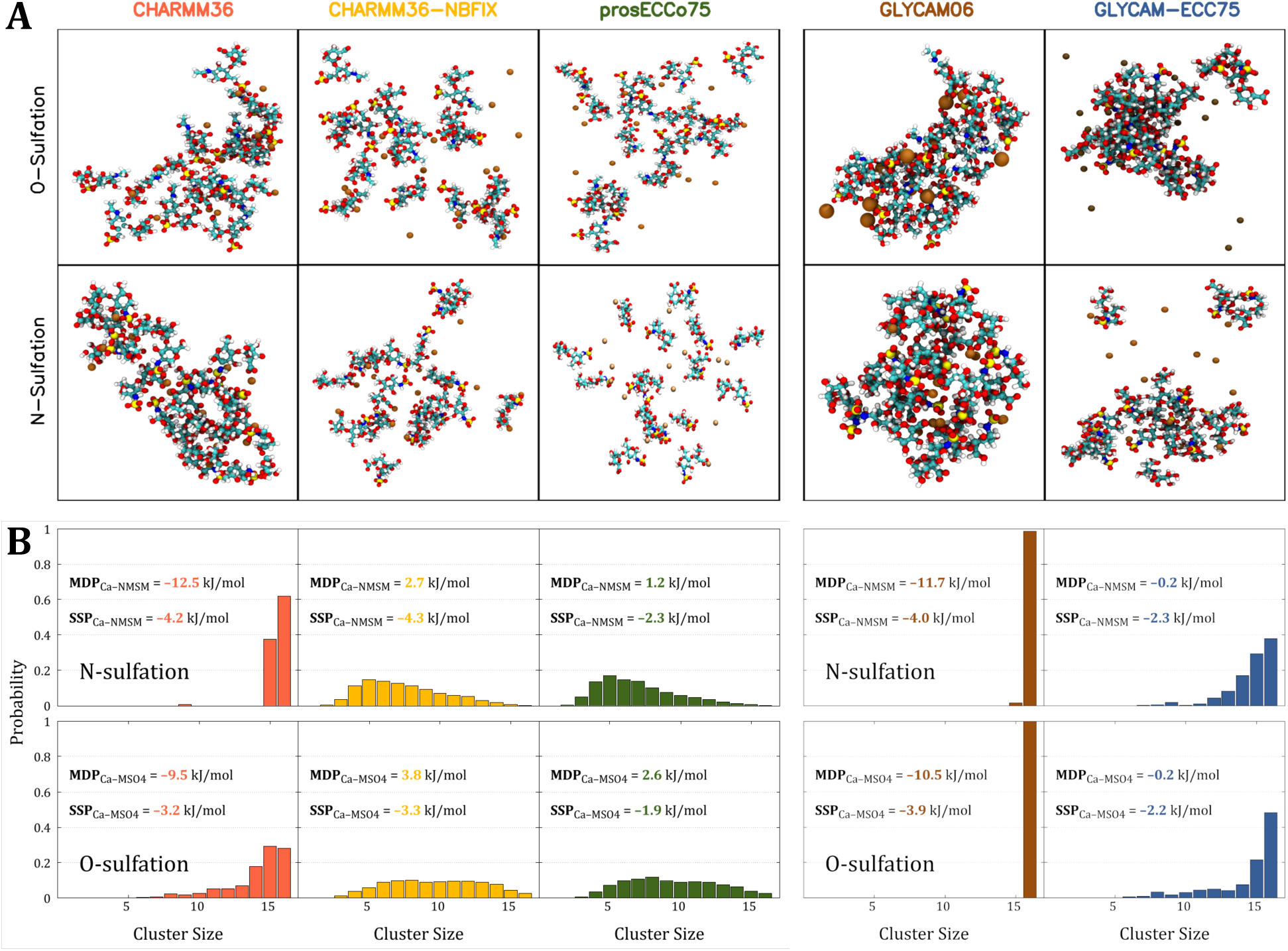
A) Representative snapshots from FFMD simulations of aqueous solutions with N- or O-sulfated GlcNAc–IdoA disaccharides and Ca^2+^. B) The cluster distribution in the FFMD simulations. The distribution shows the probability of the largest cluster having a given size — which can be as big as 16 disaccharides — observed throughout the simulation. The cluster size is defined as the number of disaccharides in continuous spatial contact with a 3.5 Å cutoff. Additionally, each panel contains absolute interaction free energies of sulfated model molecules (NMSM or MSO4) with Ca^2+^, derived from umbrella sampling simulations in a larger box with 1024 water molecules. The absolute binding energies for two interaction modes — monodentate pairing (MDP) and solvent-shared pairing (SSP) — are shown.

## Conclusions

Molecular dynamics simulations of glycosaminoglycans (GAGs) stand on the edge of significant advancements owing to increased computational capabilities and the development of more advanced models. In this work, we evaluated MD force fields for large-scale simulations of GAGs, focusing on electrostatic interactions of sulfated functional groups, which are postulated to drive GAG–GAG and GAG–protein interactions. In particular, we developed sulfated and sulfamated FF parameters for charge-scaled simulations, and compared their performance alongside other popular FFMD models. All the tested models were bench-marked against ab initio molecular dynamics (AIMD) simulations, in which we examined calcium binding to sulfated molecules that represent N- and O-sulfated motifs of GAGs. Our findings reveal a considerable variance in the performance of the tested force fields. While AIMD data consistently indicate that solvent-shared pairing is always the predominant Ca^2+^–sulfamate/sulfate interaction mode, the nonpolarizable force fields CHARMM36 and GLYCAM06, as well as the explicitly polarizable AMOEBA and Drude, overestimate contact ion pairing. In contrast, our new charge-scaled derivatives of CHARMM36 and GLYCAM06 — prosECCo75 and GLYCAM-ECC75, respectively — along with CHARMM36-NBFIX show better agreement with AIMD results, with the charge-scaled force fields clearly performing the best.

Our study further emphasizes the importance of accurate force field descriptions in simulating saccharide solutions. We observed that seemingly unimportant and often overlooked force field inaccuracies result in qualitatively distinct behaviors, *e.g.*, leading to excessive aggregation when simulating sulfated/sulfamated saccharides. Intriguingly, even models like GLYCAM-ECC75, which capture Ca^2+^ binding extremely well, still exhibit aggregation problems mainly due to imbalances in saccharide–saccharide and saccharide–water interactions. Additionally, our research demonstrates the value of employing accurate simulation models to elucidate Ca^2+^’s role as a possible modulatory agent in GAG–GAG and GAG– protein interactions.

To conclude, a critical challenge for most force fields still remains the accurate modeling of electrostatic interactions, particularly for systems with high charge density, such as those containing oxyanions and Ca^2+^. Although some simulation models can explicitly treat electronic polarizability, most force fields still inadequately represent ionic interactions. Scaling partial atomic charges by 75%, as related to the screening by the solvents’s high-frequency dielectric constant, has emerged as a promising, computationally efficient approach to ad-dress this issue. Future research should explore alternative scaling factors and potentially target the optimization of non-bonded and bonded parameters for interactions unique to sulfation. Selecting a force field that accurately captures sulfation-related interactions in GAGs is crucial for understanding their natural behavior and interactions with proteins and other biomolecules. Ultimately, accurate modeling is essential for elucidating the important role of GAGs in biology and their various applications.

## Acknowledgement

M.R.-F. and V.K. acknowledge the support from the Charles University in Prague and the International Max Planck Research School in Dresden. This work was also supported by the Ministry of Education, Youth and Sports of the Czech Republic through the e-INFRA CZ (ID:90254). We acknowledge Grammarly and ChatGPT for improving the readability and language of the manuscript.

## Supporting Information Available

Detailed description of simulation models and protocols used. Supplementary results, in-cluding additional free energy profiles.

