## Supporting Information for "Developing and Benchmarking Sulfate and Sulfamate Force Field Parameters for Glycosaminoglycans via Ab Initio Molecular Dynamics Simulations"

### Contents

**S1 Simulation Models and Implemented Adjustments to Mimic Glycosaminoglycans S-4**

S1.1 Simulation Models of Sulfated Small Molecules and Calcium . . . . . . . . S-4

S1.2 Partial Atomic Charges . . . . . . . . . . . . . . . . . . . . . . . . . . . . S-5

S1.3 Bond and Angle Parameters . . . . . . . . . . . . . . . . . . . . . . . . . S-7

S1.4 Bonds and Angles From Ab Initio Molecular Dynamics . . . . . . . . . . . S-10

**S2 Simulation Protocols S-11**

S2.1 CHARMM36 / CHARMM36–NBFIX / prosECCo75 . . . . . . . . . . . . . S-11

S2.2 GLYCAM06 / GLYCAM-ECC75 . . . . . . . . . . . . . . . . . . . . . . . S-11

S2.3 Drude . . . . . . . . . . . . . . . . . . . . . . . . . . . . . . . . . . . . . S-12

S2.4 AMOEBA . . . . . . . . . . . . . . . . . . . . . . . . . . . . . . . . . . . S-13

S2.5 Umbrella Sampling Simulations: Reference Distances and Force Constants for Harmonic Restraints . . . . . . . . . . . . . . . . . . . . . . . . . . S-14

**S3 Supplementary Results S-18**

S3.1 D3 Correction in Ab Initio Molecular Dynamics . . . . . . . . . . . . . . . S-18

S3.2 Model Molecules vs. Monosaccharides . . . . . . . . . . . . . . . . . . . . S-20

S3.3 Sampling along the Reaction Coordinate . . . . . . . . . . . . . . . . . . . S-22

S3.4 Effect of the Box Size on Free Energy Profiles . . . . . . . . . . . . . . . . S-25

S3.5 Umbrella Sampling Simulations in a Larger Box . . . . . . . . . . . . . . . S-26

S3.6 Umbrella Sampling Simulations with Adding Chloride Anion . . . . . . . . S-29

S3.7 Umbrella Sampling Simulations with vdW-modifier . . . . . . . . . . . . . S-30

S3.8 Umbrella Sampling Simulations with Larger Cutoff . . . . . . . . . . . . . S-31

S3.9 Role of Water Model on Calcium Binding . . . . . . . . . . . . . . . . . . S-32

S3.10 Accelerated Weight Histogram vs. Umbrella Sampling Simulations . . . . . S-34

S3.11 Additional Free Energy Profiles . . . . . . . . . . . . . . . . . . . . . . . S-35

S3.12 Additional Radial Distribution Functions . . . . . . . . . . . . . . . . . . . . . S-36

**References** **S-38**

### S1 Simulation Models and Implemented Adjustments to Mimic Glycosaminoglycans

#### S1.1 Simulation Models of Sulfated Small Molecules and Calcium

NMSM and MSO4 molecules were built using Ligand Modeler,[S1] implemented alongside other modules into CHARMM-GUI online utility.[S2] The CGenFF[S3,S4] was used to generate the force field (FF) parameters compatible with the most recent version of CHARMM FF.[S5] Then, we used the FF-Converter[S6] to generate the NMSM/MSO4 parameters compatible with GLYCAM06 FF.[S7,S8] The default CHARMM calcium model was used for CHARMM36 and CHARMM36–NBFIX simulations, 12-6 IOD calcium[S9] was used for GLYCAM06 simulations, while previously developed scaled-charge calcium[S10] (so-called "Ca_*s*") was used for prosECCo75 and GLYCAM-ECC75 simulations. AMOEBA parameters for the studied molecules were derived using the Poltype2 protocol,[S11] while the 2018 AMOEBA biopolymer FF was used for water and calcium cation.[S12,S13] For Drude simulations, the most recent MSO4 and NMSM parameters were adopted,[S14,S15] along with Drude-compatible SWM4-NDP water[S16] and the corresponding calcium model.[S17] Molecular coordinates were converted to Drude-compatible format using the `psfgen` plugin in VMD.[S18]

The primary objective of this study is to evaluate the accuracy of various FFs in simulating glycosaminoglycans (GAGs). To achieve this, we compared the parameters for NMSM and MSO4 molecules — specifically their sulfamate and sulfate groups — with those corresponding to GAGs. In the following discussion, we detail the critical elements of these FFs and targeted modifications applied to some of the generated parameter sets for NMSM and MSO4 to accurately resemble the sulfamate/sulfate groups found in GAGs. The related topologies and FF parameters are accessible in the Zenodo repository: 10.5281/zenodo.10036627

#### S1.2 Partial Atomic Charges

We employed CGenFF[S3,S4] and Glycan Modeler,[S19] as implemented in CHARMM-GUI,[S2] to generate CHARMM36 parameters for sulfated model molecules and saccharides, respectively. The derived partial charges are compared in Table S1. While the charges for MSO4 mirrored those in O-sulfated motifs of GAGs, those for NMSM exhibited significant discrepancies. Throughout this work, we adopted charges found in N-sulfated motifs of GAGs. Nevertheless, we checked the impact of employing CGenFF-derived charges on the free energy of calcium binding to NMSM. These parameters demonstrated inferior performance compared to those resembling GAGs, Figure S1.

Similarly, we compared the partial atomic charges for GLYCAM models. The charges for model sulfate molecules, as derived using FF-Converter,[S6] differed from those in sulfated GAGs generated by GAG builder at GLYCAM-Web (https://glycam.org/gag/), Table S2. Consequently, throughout this study, we adopted the charges used for GAGs.

No changes were made to the parameters generated for Drude and AMOEBA simulations.

**Table S1:** CHARMM36 partial atomic charges in sulfamate/sulfate groups of NMSM/MSO4 molecules and N-/O-sulfated GAGs. Values in bold and highlighted in green were used in all simulations of NMSM and MSO4 unless specified otherwise.

| **CHARMM36** | | |
|---|---|---|
| **N-sulfation** | | |
| | **NMSM** | **GAGs** |
| Sulfur | 0.62* | **1.11** |
| Sulfamate oxygen | –0.50* | **–0.64** |
| Amine nitrogen | –0.52* | **–0.73** |
| Amine hydrogen | 0.33* | **0.35** |
| **O-sulfation** | | |
| | **MSO4** | **GAGs** |
| Sulfur | **1.33** | 1.33 |
| Sulfate oxygen | **–0.65** | –0.65 |
| Bridging oxygen | **–0.28** | –0.28 |

*) Parameter is different compared to GAGs

**Table S2:** GLYCAM06 partial atomic charges in sulfamate/sulfate groups of NMSM/MSO4 molecules and N-/O-sulfated GAGs. Values in bold and highlighted in green were used in all simulations of NMSM and MSO4 unless specified otherwise.

| | **GLYCAM06** | |
|---|---|---|
| | **N-sulfation** | |
| | **NMSM** | **GAGs** |
| Sulfur | 1.531867* | **1.245** |
| Sulfamate oxygen | –0.7519* | **–0.694** |
| Amine nitrogen | –0.9181* | **–0.643** |
| Amine hydrogen | 0.3958* | **0.236** |
| | **O-sulfation** | |
| | **MSO4** | **GAGs** |
| Sulfur | 1.6943* | **1.245** |
| Sulfate oxygen | –0.757533* | **–0.694** |
| Bridging oxygen | –0.6085* | **–0.430** |

*) Parameter is different compared to GAGs

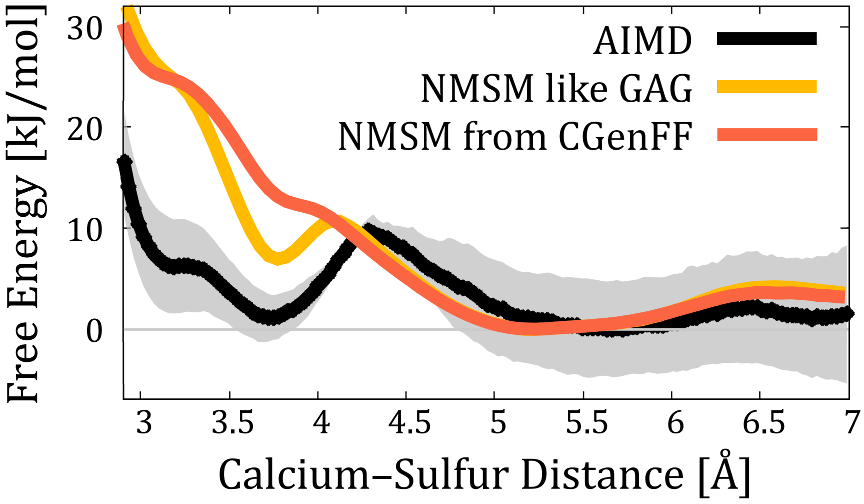


**Figure S1:** Interaction free energy profiles obtained from umbrella sampling simulations of calcium binding to NMSM using CHARMM-NBFIX FF in combination with the partial charges either adopted from sulfamate groups in GAGs (all simulations in the main text) or generated by CGenFF, see Table S1. The energy profiles are compared to those from AIMD simulations.

#### S1.3 Bond and Angle Parameters

The bond and angle parameters for sulfamate/sulfate groups within CHARMM36/prosECCo75 and GLYCAM06/GLYCAM-ECC75 FFs are given in Tables S3, S4, S5, and S6. Note that CHARMM36 and prosECCo75, as well as GLYCAM06 and GLYCAM-ECC75, share identical bond and angle parameters. Parameters that are bold and highlighted in green were used in all simulations of NMSM and MSO4 unless specified otherwise. Modifications were made to align the NMSM/MSO4 parameterization more closely with sulfamate/sulfate groups in GAGs. For CHARMM36/prosECCo75, only a single alteration was necessary from the original CGenFF parameters to ensure the geometrical stability of NMSM molecule, see Figure S2. For GLYCAM06/GLYCAM-ECC75, the parameters for NMSM and MSO4 were directly replicated from those of sulfamate/sulfate groups in GAGs due to the significant disparities with parameters generated by the FF-converter.

**Table S3:** CHARMM36/prosECCo75 bond parameters of NMSM/MSO4 molecules and corresponding sulfamate/sulfate groups in GAGs. Values in bold and highlighted in green were used in all simulations of NMSM and MSO4 unless specified otherwise. The functional form of the harmonic bond potential used in GROMACS is implied.

| **CHARMM36/prosECCo75** | | | | |
|---|---|---|---|---|
| Bond | $b_0$ [nm] | $k_b$ [kJ·mol$^{-1}$·nm$^{-2}$] | $b_0$ [nm] | $k_b$ [kJ·mol$^{-1}$·nm$^{-2}$] |
| **N-sulfation** | | | | |
| | NMSM | | GAGs | |
| S–$O_S$ | **0.1448** | **451872** | 0.1448 | 451872 |
| S–N | **0.1700** | **187443.2*** | 0.1700 | 154808 |
| **O-sulfation** | | | | |
| | MSO4 | | GAGs | |
| S–$O_S$ | **0.1448** | **451872** | 0.1448 | 451872 |
| S–$O_b$ | **0.1575*** | **209200** | 0.1610 | 209200 |

*) Parameter is different compared to GAGs

**Table S4:** CHARMM36/prosECCo75 angle parameters of NMSM/MSO4 molecules and corresponding sulfamate/sulfate groups in GAGs. Values in bold and highlighted in green were used in all simulations of NMSM and MSO4 unless specified otherwise. The functional form of the harmonic angle potential used in GROMACS is implied.

| **CHARMM36/prosECCo75** | | | | |
|---|---|---|---|---|
| Angle | $\theta_0$ [deg] | $k_\theta$ [kJ·mol$^{-1}$·rad$^{-2}$] | $\theta_0$ [deg] | $k_\theta$ [kJ·mol$^{-1}$·rad$^{-2}$] |
| **N-sulfation** | | | | |
| | NMSM | | GAGs | |
| $O_S$–S–$O_S$ | **109.47** | **1087.84** | 109.47 | 1087.84 |
| $O_S$–S–N | 94.2* | 277.8176* | **103.00** | **711.28** |
| **O-sulfation** | | | | |
| | MSO4 | | GAGs | |
| $O_S$–S–$O_S$ | **109.47** | **1087.84** | 109.47 | 1087.84 |
| $O_S$–S–$O_b$ | **98.00** | **711.28** | 98.00 | 711.28 |

*) Parameter is different compared to GAGs

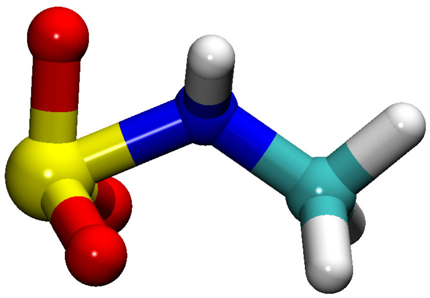

**Figure S2:** NMSM molecule after a simulation with CGenFF-generated CHARMM36 parameters. To prevent unrealistic orientation of sulfamate oxygens, the reference $O_S$–S–N angle was increased from 94.2 to 103 degrees, while the force constant of the corresponding harmonic potential was increased from 277.8176 to 711.28 kJ·mol$^{-1}$·rad$^{-2}$, Table S4.

**Table S5:** GLYCAM06/GLYCAM-ECC75 bond parameters of NMSM/MSO4 molecules and corresponding sulfamate/sulfate groups in GAGs. Values in bold and highlighted in green were used in all simulations of NMSM and MSO4 unless specified otherwise. The functional form of the harmonic bond potential used in GROMACS is implied.

| GLYCAM06/GLYCAM-ECC75 | | | | |
|---|---|---|---|---|
| Bond | $b_0$ [nm] | $k_b$ [kJ·mol$^{-1}$·nm$^{-2}$] | $b_0$ [nm] | $k_b$ [kJ·mol$^{-1}$·nm$^{-2}$] |
| **N-sulfation** | | | | |
| | NMSM | | GAGs | |
| S–$O_S$ | 0.1453* | 571534.4* | **0.1440** | **518820** |
| S–N | 0.1672* | 265349.3* | **0.1675** | **199160** |
| **O-sulfation** | | | | |
| | MSO4 | | GAGs | |
| S–$O_S$ | 0.1453* | 571534.4* | **0.1440** | **518820** |
| S–$O_b$ | 0.1623* | 324511.0* | **0.1589** | **172380** |

*) Parameter is different compared to GAGs

**Table S6:** GLYCAM06/GLYCAM-ECC75 angle parameters of NMSM/MSO4 molecules and corresponding sulfamate/sulfate groups in GAGs. Values in bold and highlighted in green were used in all simulations of NMSM and MSO4 unless specified otherwise. The functional form of the harmonic angle potential used in GROMACS is implied.

| GLYCAM06/GLYCAM-ECC75 | | | | |
|---|---|---|---|---|
| Angle | $\theta_0$ [deg] | $k_\theta$ [kJ·mol$^{-1}$·rad$^{-2}$] | $\theta_0$ [deg] | $k_\theta$ [kJ·mol$^{-1}$·rad$^{-2}$] |
| **N-sulfation** | | | | |
| | NMSM | | GAGs | |
| $O_S$–S–$O_S$ | 120.05* | 1072.7776* | **113.91** | **1029.30** |
| $O_S$–S–N | 107.43* | 595.8016* | **108.00** | **702.91** |
| **O-sulfation** | | | | |
| | MSO4 | | GAGs | |
| $O_S$–S–$O_S$ | 120.05* | 1072.7776* | **113.91** | **1029.30** |
| $O_S$–S–$O_b$ | 108.56* | 1059.3888* | **106.87** | **870.27** |

*) Parameter is different compared to GAGs

#### S1.4 Bonds and Angles From Ab Initio Molecular Dynamics

While our study does not aim to adjust the bond and angle parameters of the studied force fields based on ab initio molecular dynamics (AIMD) simulations, we did assess their comparability. Table S7 summarizes all relevant bonds and angles calculated from AIMD. Overall, we find the parameters available in CHARMM and GLYCAM models, c.f. Tables S3, S4, S5, and S6, comparable to those derived from the AIMD simulations.

**Table S7:** Average bonds and angles in sulfamate/sulfate groups of NMSM/MSO4 molecules from AIMD simulations. The data were calculated from the umbrella sampling window where the calcium cation and the studied molecules are at their maximum distance. Error estimates are calculated through block averaging using `gmx analyze` tool in GROMACS.

| **Ab Initio Molecular Dynamics** | | |
|---|---|---|
| **Bonds** | | |
| Bond | $b_0$ [nm] | |
| | NMSM | MSO4 |
| S–$O_S$ | 0.149 ± 0.0 | 0.148 ± 0.0 |
| S–N | 0.171 ± 0.0 | – |
| S–$O_b$ | – | 0.166 ± 0.0 |
| **Angles** | | |
| Angle | $\theta_0$ [deg] | |
| | NMSM | MSO4 |
| $O_S$–S–$O_S$ | 112.8 ± 0.1 | 113.7 ± 0.0 |
| $O_S$–S–N | 105.8 ± 0.1 | – |
| $O_S$–S–$O_b$ | – | 104.7 ± 0.0 |

### S2 Simulation Protocols

#### S2.1 CHARMM36 / CHARMM36–NBFIX / prosECCo75

All simulations with CHARMM36, CHARMM36–NBFIX, and prosECCo75 FFs were performed using a leap-frog integrator with a time step of 2 fs in GROMACS simulation engine.[S20] Buffered Verlet lists[S21] were used to keep track of atomic neighbors. Temperature of 300 K was maintained using the Nosé–Hoover thermostat[S22,S23] with a 1 ps coupling time. In $NPT$ simulations, pressure was kept at 1 bar via the Parrinello-Rahman barostat[S24] with a relaxation time of 5 ps and compressibility of $4.5\times10^{-5}$ bar$^{-1}$. Electrostatic interactions were computed with a cutoff set to 1.2 nm, while the smooth particle mesh Ewald method was used to treat the long-range electrostatics.[S25] van der Waals interactions were computed with a cutoff of 1.2 nm and force switching function at 1.0 nm.[S26] For umbrella sampling simulations in a small box containing only 128 water molecules, the cutoff for both electrostatic and Lennard Jones interactions was reduced to 0.7 nm, without modifications to the van der Waals potential. LINCS algorithm was used to constraint heavy atom–hydrogen bonds.[S27,S28] Water was simulated with a rigid CHARMM-specific TIP3P model[S29,S30] constrained using SETTLE.[S31]

#### S2.2 GLYCAM06 / GLYCAM-ECC75

All simulations with GLYCAM06 and GLYCAM-ECC75 FFs were performed using a leap-frog integrator with a time step of 2 fs in GROMACS simulation engine.[S20] Buffered Verlet lists[S21] were used to keep track of atomic neighbors. Temperature of 300 K was maintained using the Nosé–Hoover thermostat[S22,S23] with a 1 ps coupling time. In $NPT$ simulations, pressure was kept at 1 bar via the Parrinello-Rahman barostat[S24] with a relaxation time of 5 ps and compressibility of $4.5\times10^{-5}$ bar$^{-1}$. The long-range dispersion correction for energy and pressure was applied. Electrostatic interactions were computed with a cutoff set to 1.0 nm, while the smooth particle mesh Ewald method was used to treat the long-range electrostatics.[S25] Lennard–Jones interactions were cut off of 1.0 nm by shifting the van der Waals potential by a constant. For umbrella sampling simulations in a small box containing only 128 water molecules, the cutoff for both electrostatic and Lennard–Jones interactions was reduced to 0.7 nm. LINCS algorithm was used to constraint heavy atom–hydrogen bonds.[S27,S28] Water was simulated with a rigid TIP3P model[S29] constrained using SETTLE.[S31] Note that unlike in other Amber force fields, GLYCAM FFs do not apply any scaling to 1–4 interactions.[S7]

#### S2.3 Drude

Drude simulations were performed using the NAMD package (nightly build as of 15th of July 2022).[S32] The `colvars` module[S33] was used for restraining calcium–sulfur distances in umbrella sampling simulations. Each umbrella sampling window was minimized for 5000 steps, equilibrated for 1 ns, and then simulated with a 1 fs time step. The temperature was kept at 300 K using a Langevin thermostat[S34] with a 1 $ps^{-1}$ damping. The temperature of the Drude particles was maintained at 1 K using a separate Langevin thermostat with 20 $ps^{-1}$ damping. The Drude bond length was kept under 0.2 Å with a hard-wall constraint.[S35] The non-bonded Thole screening (NBTHOLE) was turned on with a default radius of 5 Å.

During Drude umbrella sampling simulations, we encountered problems with overpolarization,[S17] which is a well-recognized issue common for highly polarizable ions. In that case, the interactions between negative and positive moieties can be so strong that the Drude particles are displaced too far from their parent atoms, and the molecular geometry is compromised. A few approaches to palliate this problem are possible, with the most prominent being employing NBFIX or NBTHOLE parameters to dampen the exaggerated interactions.[S36] As fitting Drude FF is outside the scope of this work, here we report the interaction free energies of $Ca^{2+}$ binding to MSO4/NMSM using the available parameters.[S14]

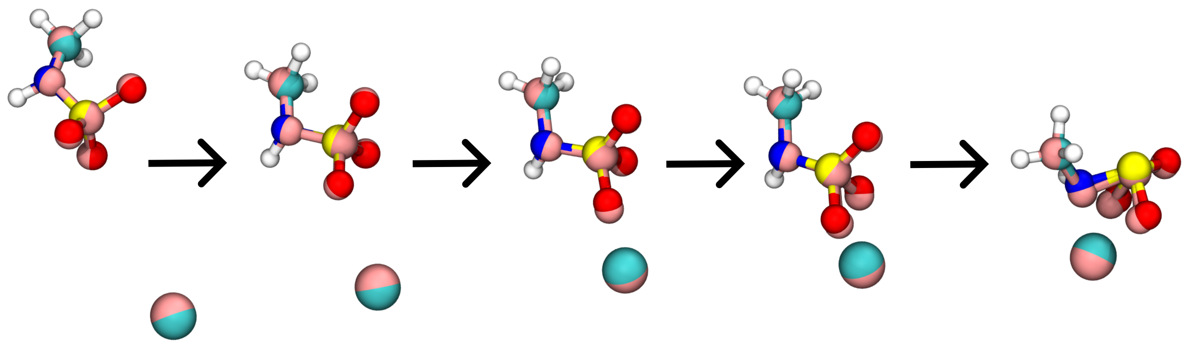

**Figure S3:** Time evolution of the binding between $Ca^{2+}$ and NMSM. The Drude particles are shown as the pink spheres surrounding most atoms. The breakage of the NMSM geometry can be observed as the simulation proceeds.

#### S2.4 AMOEBA

AMOEBA simulations were performed using Tinker9 software (https://github.com/TinkerTools/tinker9). The r-RESPA multiple-time-step integration scheme[S37] was employed, with 2 fs and 0.25 fs time steps for the nonbonded and bonded interactions, respectively. Temperature of 300 K was controlled by stochastic velocity rescaling thermostat[S38] with a time constant of 0.2 ps.

As mentioned earlier, we used Poltype2 protocol[S11] to generate parameters for NMSM and MSO4 molecules. We also tested the earlier version of the protocol, Poltype.[S39] The results obtained with the original parameterization scheme were found significantly worse than those using Poltype2 (data not shown).

#### S2.5 Umbrella Sampling Simulations: Reference Distances and Force Constants for Harmonic Restraints

**Table S8:** The reference distances ($r_{\text{ref}}$) and force constants ($k$) of the harmonic restraints in the AIMD and FFMD umbrella sampling simulations (except for simulations with Drude, see below). The larger force constants at larger distances were required to keep the ions at a certain distance without taking into account periodic boundary conditions due to specific implementation of umbrella sampling simulations in GROMACS software. The functional form of the harmonic potential used in GROMACS is implied. Note that, in case of AMOEBA simulations, the "input" force constant of the harmonic potential should be smaller by a factor of two due to the missing scaling factor in the equation of the harmonic potential in Tinker software.

| Reference distance, $r_{\text{ref}}$ [nm] | Force constant, $k$ (MSO4 / NMSM) [kJ·mol$^{-1}$·nm$^{-2}$] |
|---|---|
| 0.28 | 20000 / 10000 |
| 0.30 | 20000 / 10000 |
| 0.32 | 20000 / 10000 |
| 0.34 | 20000 / 10000 |
| 0.36 | 20000 / 10000 |
| 0.38 | 20000 / 10000 |
| 0.40 | 20000 / 10000 |
| 0.42 | 20000 / 10000 |
| 0.44 | 20000 / 10000 |
| 0.46 | 5000 / 2500 |
| 0.50 | 2500 / 2500 |
| 0.54 | 2500 / 2500 |
| 0.58 | 2500 / 2500 |
| 0.62 | 2500* / 2500 |
| 0.66 | 5000 / 5000 |

*) For some force fields, the force constant was increased to 5000 kJ·mol$^{-1}$·nm$^{-2}$

**Table S9:** The reference distances ($r_{\mathrm{ref}}$) and force constants ($k$) of the harmonic restraints in the umbrella sampling simulations of MSO4 with Drude force field. The addition of extra windows and a notable increase of most force constants was needed to keep the distances at the reference values and ensure windows overlapping. Please note that the force constants and distances are given in NAMD units, Å and kcal·mol$^{-1}$·Å$^{-2}$.

| Reference distance, $r_{\mathrm{ref}}$ [nm] | Force constant, $k$ [kcal·mol$^{-1}$·Å$^{-2}$] |
|---|---|
| 0.295 | 334.6 |
| 0.32 | 95.6 |
| 0.34 | 95.6 |
| 0.36 | 95.6 |
| 0.38 | 95.6 |
| 0.39 | 286.8 |
| 0.40 | 143.4 |
| 0.41 | 286.8 |
| 0.42 | 143.4 |
| 0.44 | 95.6 |
| 0.46 | 47.8 |
| 0.50 | 5.98 |
| 0.54 | 5.98 |
| 0.58 | 5.98 |
| 0.62 | 5.98 |
| 0.66 | 11.94 |

**Table S10:** The reference distances ($r_{\text{ref}}$) and force constants ($k$) of the harmonic potential in the umbrella sampling simulations of NMSM with Drude force field. The addition of extra windows and a notable increase of most force constants was needed to overcome the high forces between NMSM and calcium in Drude. All the windows were prepared and simulated, however none of the distances shorter than 0.54 nm was finished due to simulation errors caused by the overpolarization of NMSM. Please note that the force constants and distances are given in NAMD units, Å and [kcal·mol$^{-1}$·Å$^{-2}$].

| Reference distance, $r_{\text{ref}}$ [nm] | Force constant, $k$ [kcal·mol$^{-1}$·Å$^{-2}$] |
|---|---|
| 0.28 | 800.65 |
| 0.30 | 800.65 |
| 0.32 | 800.65 |
| 0.34 | 800.65 |
| 0.36 | 537.75 |
| 0.38 | 800.65 |
| 0.40 | 800.65 |
| 0.42 | 800.65 |
| 0.44 | 800.65 |
| 0.46 | 2.99 |
| 0.50 | 2.99 |
| 0.54 | 2.99 |
| 0.58 | 2.99 |
| 0.62 | 2.99 |
| 0.66 | 2.99 |

**Table S11:** The reference distances ($r_{\text{ref}}$) and force constants ($k$) of the harmonic potential in the FFMD umbrella sampling simulations in a large simulation box size. The values for shorter distances, if not explicitly given here, are the same as in Table S8. The larger force constants at larger distances were required to keep the ions at a certain distance without taking into account periodic boundary conditions due to the specific implementation of umbrella sampling simulations in GROMACS software. The functional form of the harmonic potential used in GROMACS is implied.

| Reference distance, $r_{\text{ref}}$ [nm] | Force constant, $k$ [kJ·mol$^{-1}$·nm$^{-2}$] |
|---|---|
| 0.66 | 2500 |
| 0.70 | 1000 |
| 0.75 | 1000 |
| 0.80 | 1000 |
| 0.85 | 1000 |
| 0.90 | 1000 |
| 0.95 | 1000 |
| 1.00 | 1000 |
| 1.05 | 1000 |
| 1.10 | 1000 |
| 1.15 | 1000 |
| 1.20 | 1000 |
| 1.25 | 1000 |
| 1.30 | 1000* |
| 1.35 | 2500 |
| 1.40 | 5000 |

*) For some force fields, the force constant was increased to 2500 kJ·mol$^{-1}$·nm$^{-2}$

### S3 Supplementary Results

To verify the viability of our methodology, particularly umbrella sampling simulations, and to further explore the topic, we conducted multiple checks and additional tests on selected systems.

#### S3.1 D3 Correction in Ab Initio Molecular Dynamics

A recent work[S40] showed that commonly used in AIMD simulations D3 correction[S41] should not be applied to alkali and alkali earth cations. Our data, Figure S4, compare the free energy profiles calculated from AIMD simulations without (used throughout our work) and with D3 correction applied to calcium. We observe significant differences between the profiles, thereby highlighting that improper protocol in using D3 correction can substantially alter the AIMD results and following conclusions.

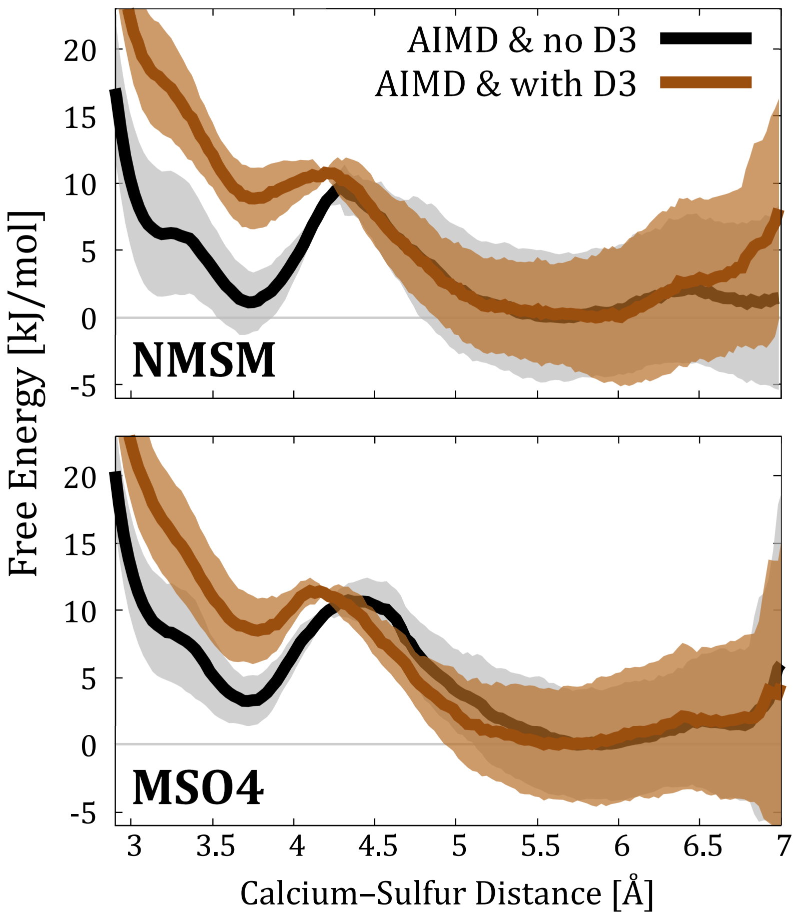


**Figure S4:** The AIMD interaction free energy profiles obtained with and without D3 correction applied to calcium cation.

#### S3.2 Model Molecules vs. Monosaccharides

We performed umbrella sampling simulations of calcium binding to sulfated monosaccharides, Figure S5, to check whether the free energy profiles are consistent with those for model molecules. Our results, Figure S6, show that the free energy profiles are essentially identical.

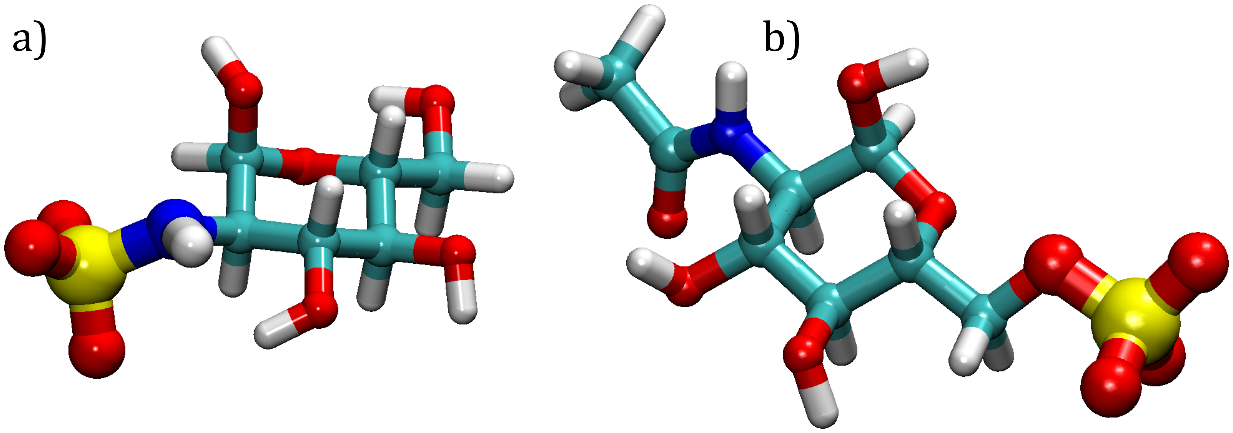


**Figure S5:** Tested monosaccharides: (a) N-sulfated N-acetyl-D-glucosamine and (b) 6-O-sulfated N-acetyl-D-glucosamine.

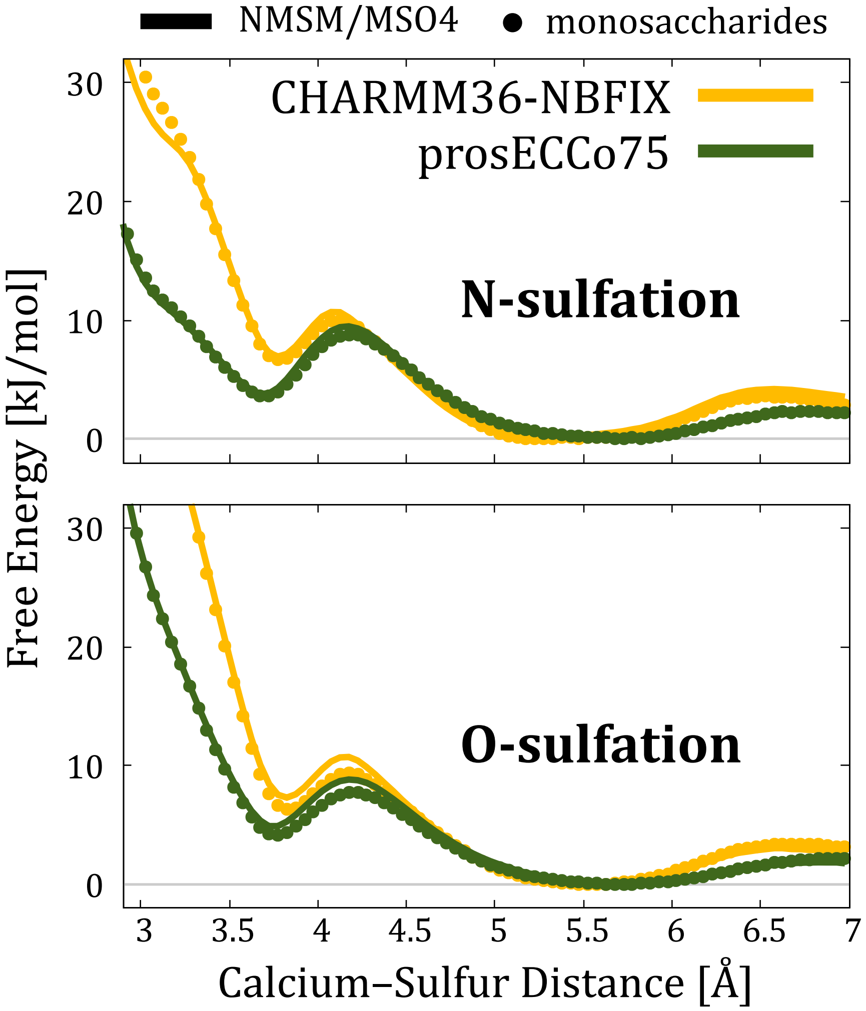


**Figure S6:** Interaction free energy profiles obtained from FFMD umbrella sampling simulations of calcium binding to NMSM and MSO4 molecules (solid lines) in comparison to those of calcium binding to N- and O-sulfated monosaccharides (points), Figure S5. Due to larger size of monosaccharides, the umbrella sampling simulations with the monosaccharides were performed in a larger simulation box with 1024 water molecules. In this comparison, all energy profiles were shifted to zero in the position of the solvent-shared pairing minimum.

#### S3.3 Sampling along the Reaction Coordinate

For simplicity and consistency, in all our simulations, the collective variable was the distance between calcium cation and sulfur atom of either MSO4 or NMSM. We performed additional simulations with the collective variable being the distance between calcium and the center of mass (COM) of the corresponding anion. Our results, Figure S7, show that the free energy profiles are essentially the same except a little offset related to the shifted location of the COM with respect to sulfur atom. Therefore, our initial choice of the collective variable is suitable and does not preclude any interaction motifs.

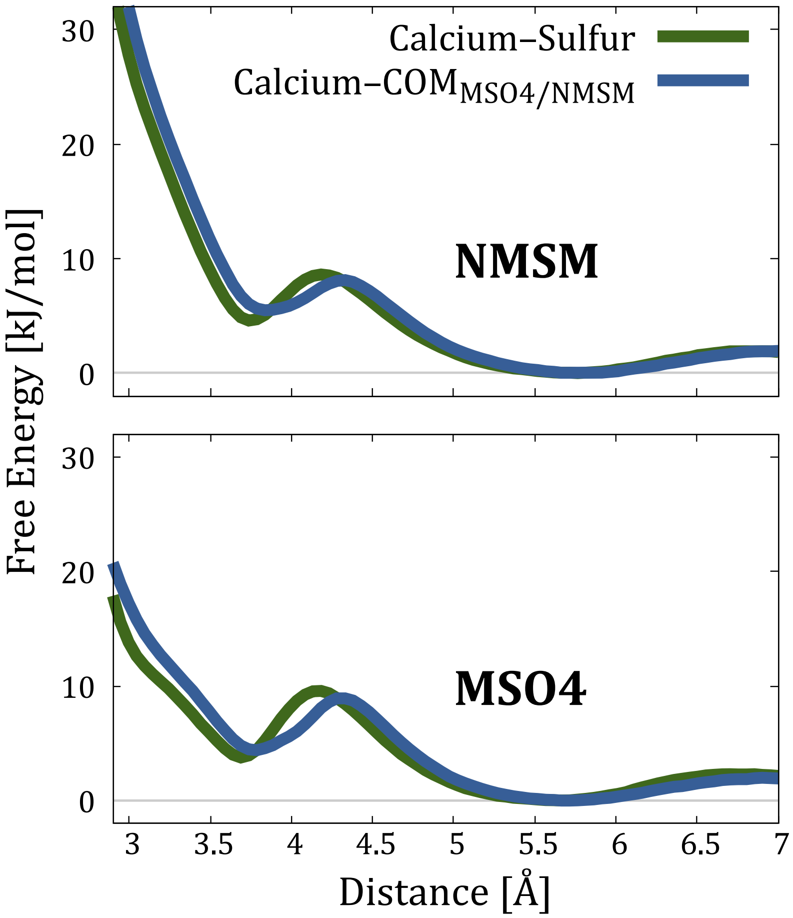


**Figure S7:** Interaction free energy profiles obtained from accelerated weight histogram (AWH) simulations of calcium binding to MSO4 using prosECCo75 force field and different interatomic/intermolecular distances as the collective variable.

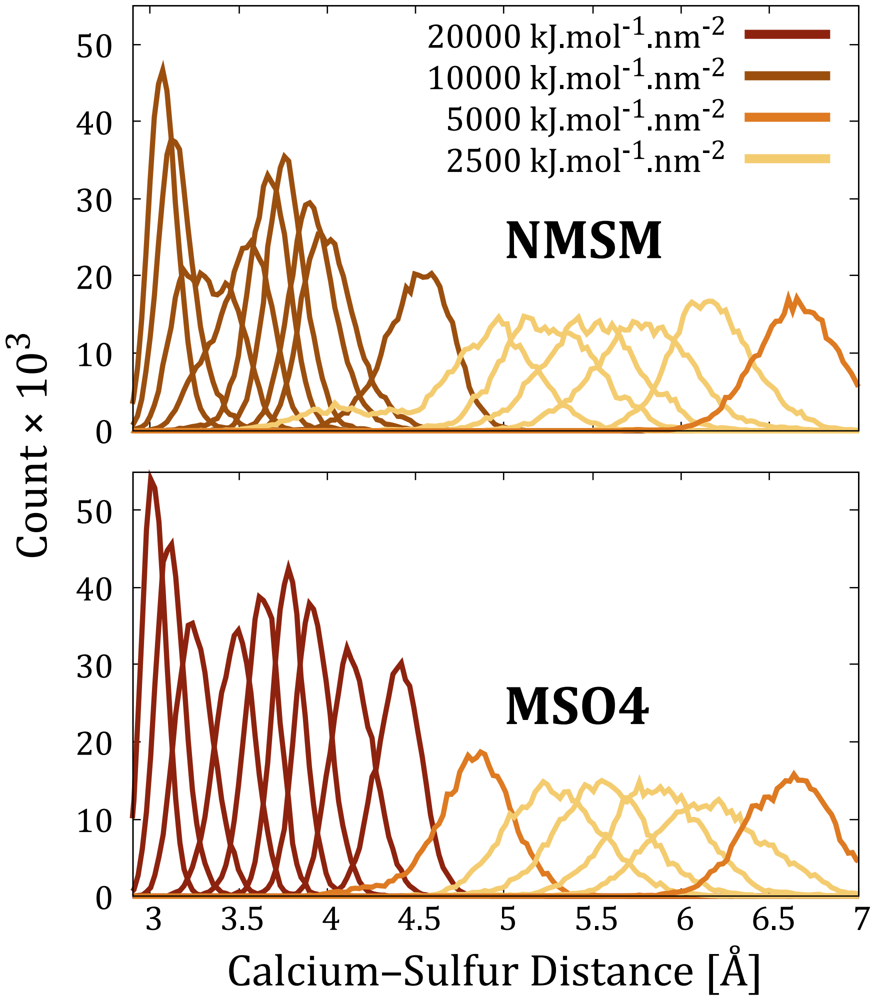


**Figure S8:** The sampling histogram along the reaction coordinate from the AIMD umbrella sampling simulations.

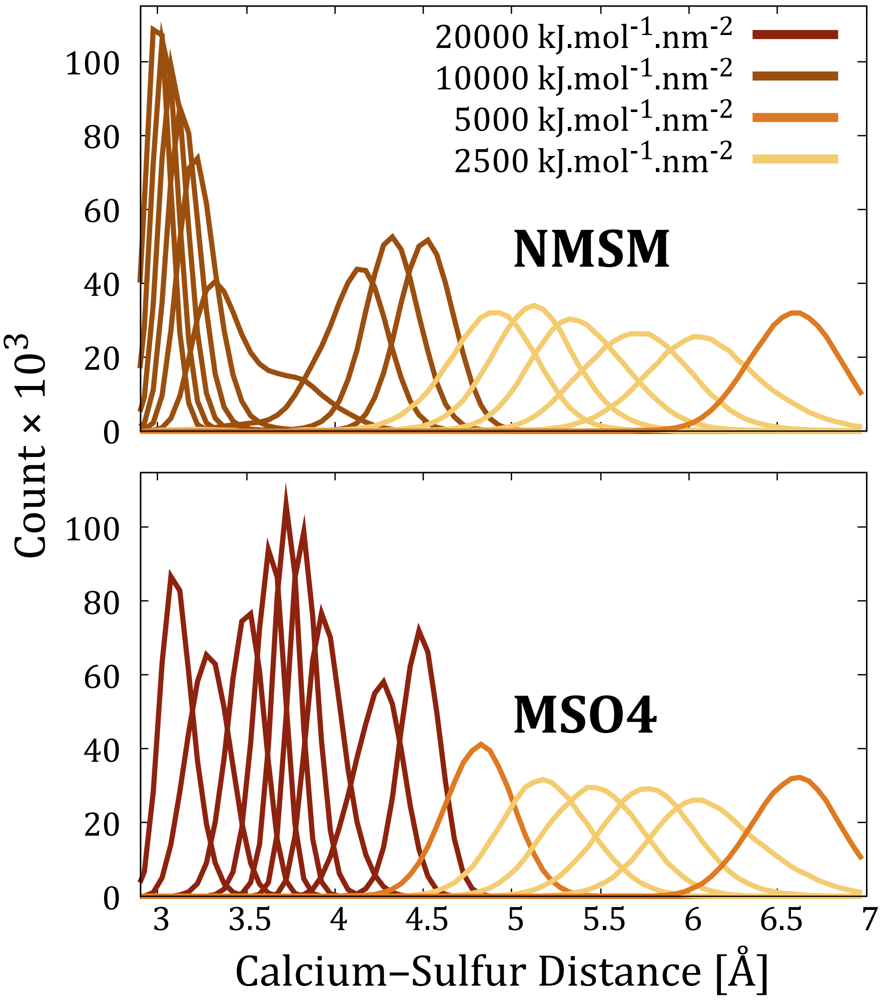


**Figure S9:** An example sampling histogram along the reaction coordinate from the FFMD umbrella sampling simulations, namely from the simulations with CHARMM36-NBFIX force field.

#### S3.4 Effect of the Box Size on Free Energy Profiles

In both AIMD and FFMD umbrella sampling simulations, the box size was fixed (*NVT* ensemble) for simulations with a given molecule (MSO4 and NMSM) and selected from *NPT* simulations with prosECCo75 force field. Despite the fact that minor deviations in the box size were present in simulations with different force fields when performed in *NPT* ensemble (as we tested some of them, the size of smaller box was in a range between 15.8 and 16 Å), our results, Figure S10, show that the impact of the box size on the free energy profiles is essentially zero.

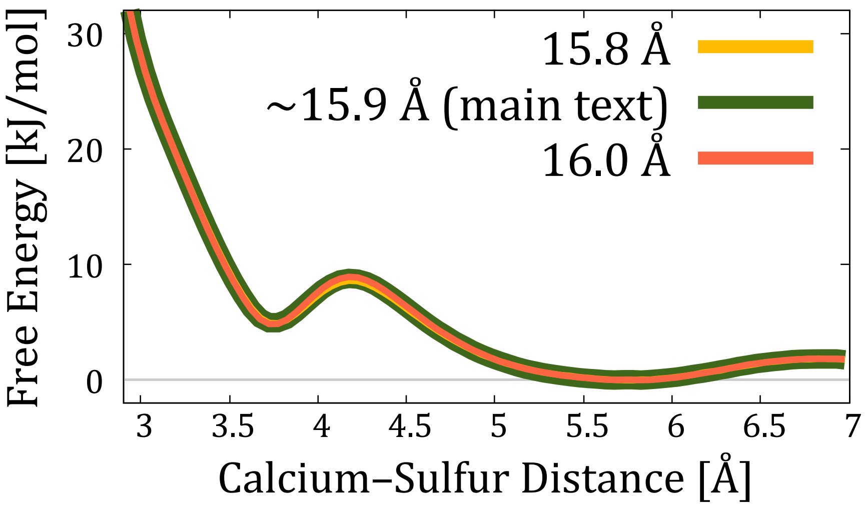


**Figure S10:** Interaction free energy profiles obtained from umbrella sampling simulations of calcium binding to MSO4 using prosECCo75 force field, varying the fixed box size in *NVT* ensemble.

#### S3.5 Umbrella Sampling Simulations in a Larger Box

In order to estimate the absolute binding free energy of calcium binding, not only the difference between contact and solvent-shared ion pairing, we performed additional simulations with selected force fields in a larger simulation box with 1024 water molecules.

First, we compared whether the energy profiles match when both were shifted to zero in the position of the solvent-shared pairing minimum, Figure S11. Then, we shifted the profiles from the simulations in a larger box to zero in the region of the bulk, Figure S12. In this case, we could estimate the absolute free energies of binding for different force fields as summarized in Figure 6.

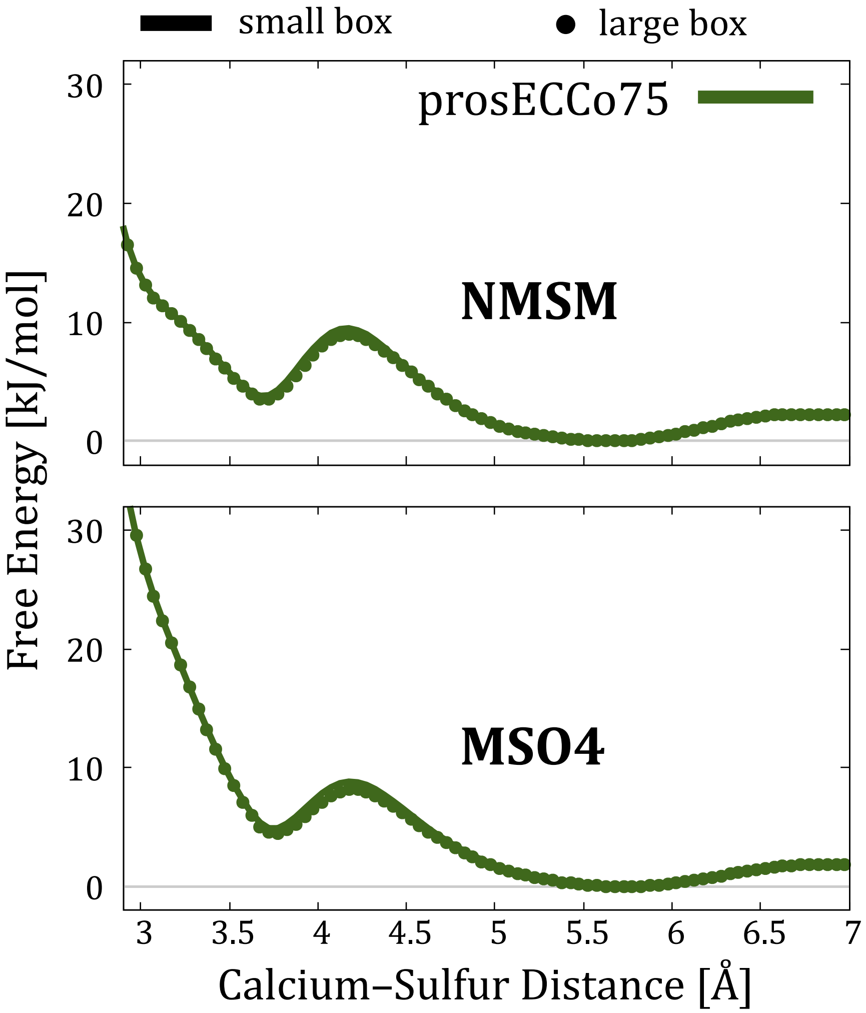


**Figure S11:** Comparison of interaction free energy profiles obtained from umbrella sampling simulations in a small (128 water molecules, used in most simulations, including AIMD) and large (1024 water molecules) simulation boxes.

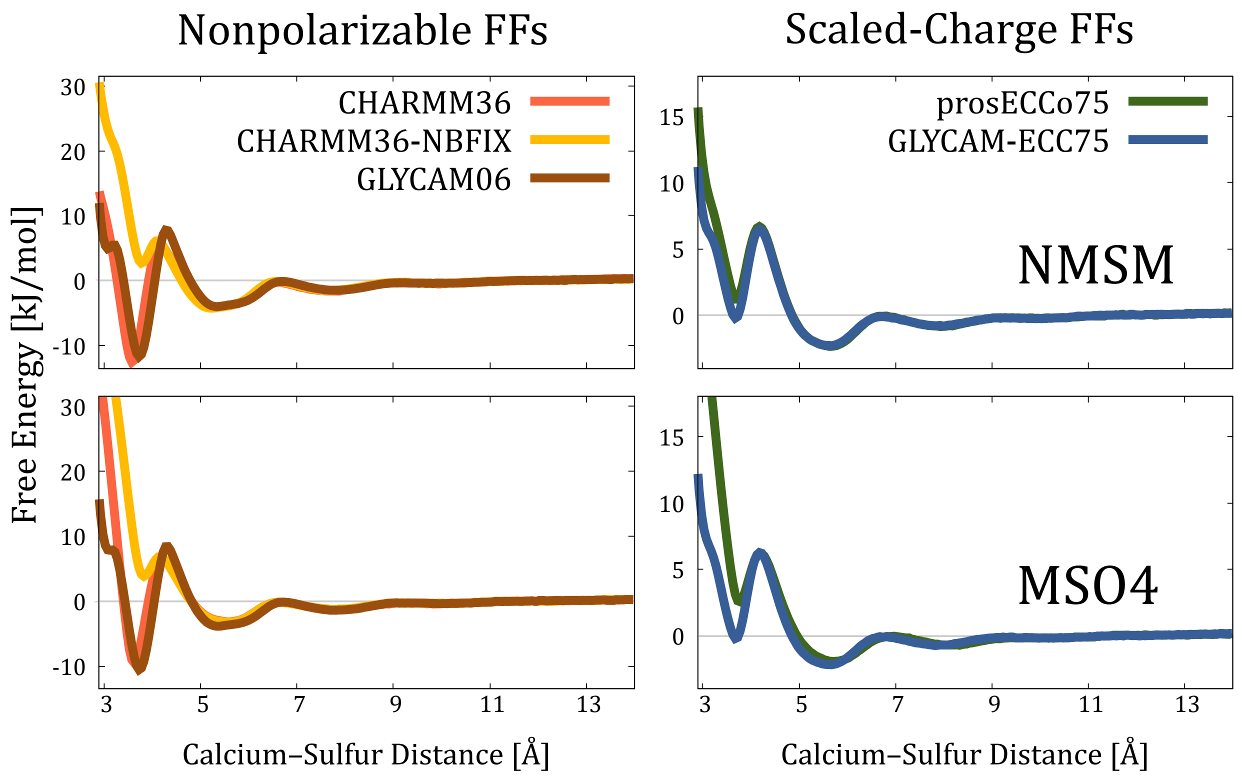


**Figure S12:** Interaction free energy profiles of calcium binding to MSO4 and NMSM molecules obtained from umbrella sampling simulations in a larger box.

#### S3.6 Umbrella Sampling Simulations with Adding Chloride Anion

Since the system with calcium cation and either MSO4 or NMSM anion is not charge-neutral (+2 vs. −1), the remaining positive charge is neutralized by a background charge. One may argue that this may affect the electrostatic interactions in a relatively small box. To eliminate this possibility, we performed FFMD umbrella sampling simulations with $Cl^-$ anion added to the system. Our results, Figure S13, show that there is a zero difference between these two approaches.

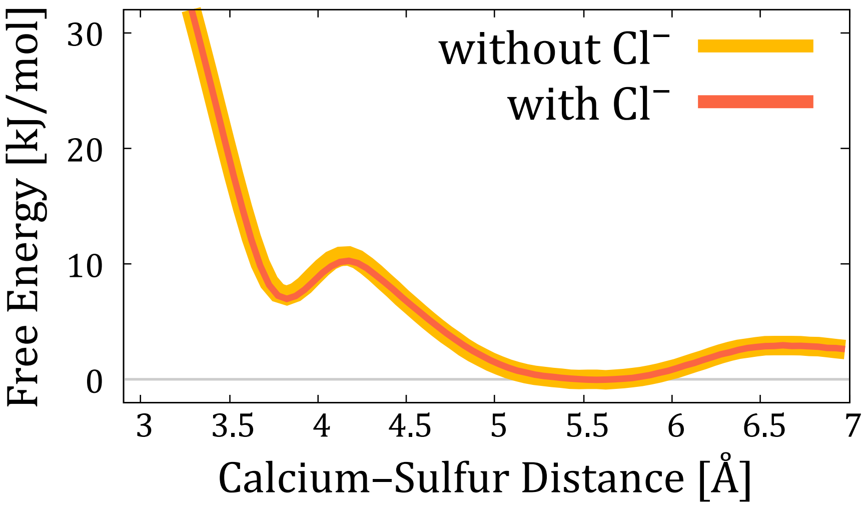


**Figure S13:** Interaction free energy profiles obtained from umbrella sampling simulations of calcium binding to MSO4 using CHARMM36-NBFIX model without (as in all results presented in the main text) and chloride ion present in the system.

##### S3.7 Umbrella Sampling Simulations with vdW-modifier

The recommended protocol for MD simulations with CHARMM36 (and, therefore, prosECCo75) force field is using van der Waals cutoff scheme with the cutoff value set to 1.2 nm and forces smoothly switched to zero starting at 1.0 nm, which can be set in GROMACS by `vdw-modifier = Force-switch`. Given the half box size in our simulations is ∼0.8 nm, we did not use any van der Waals switch, which can be set in GROMACS by `vdw-modifier = None`, while the cutoff was set to 0.7 nm. The same cutoff was applied to electrostatic interactions. To check whether there is any effect of the van der Waals treatment on our data, we performed umbrella sampling simulations with `vdw-modifier = Potential-shift`, which is the default option in GROMACS and the one used in all GLYCAM simulations. Our results, Figure S14, show that there is a zero difference between these two choices.

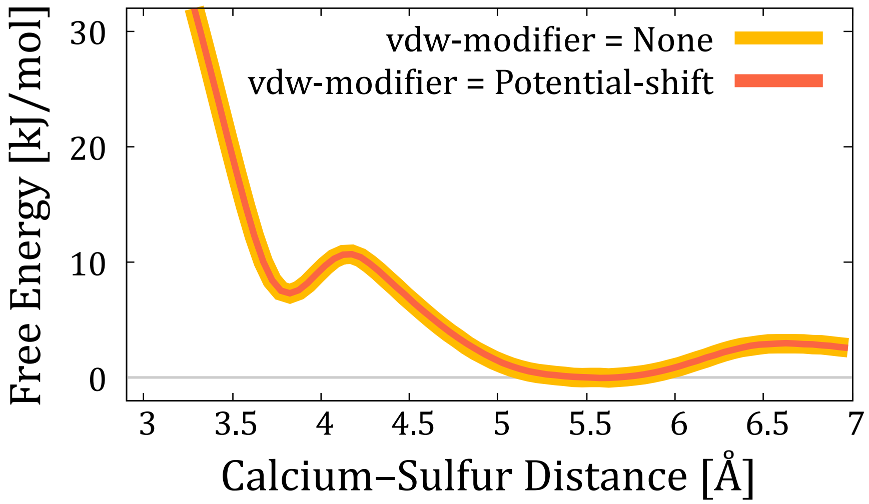


**Figure S14:** Interaction free energy profiles obtained from umbrella sampling simulations of calcium binding to MSO4 using CHARMM36-NBFIX model with vdw-modifier set to "None" (orange line, used in all CHARMM36 and prosECCo75 simulations in the main text) or "Potential-shift" (red line, used in all GLYCAM06 and GLYCAM-ECC75 simulations in the main text).

#### S3.8 Umbrella Sampling Simulations with Larger Cutoff

Similarly, we tested if the overall change in the cutoff from 0.7 nm to 0.75 nm (for both van der Waals and electrostatic interactions) has any impact on the free energy profiles. Our results, Figure S15, show that the corresponding free energy profiles are identical.

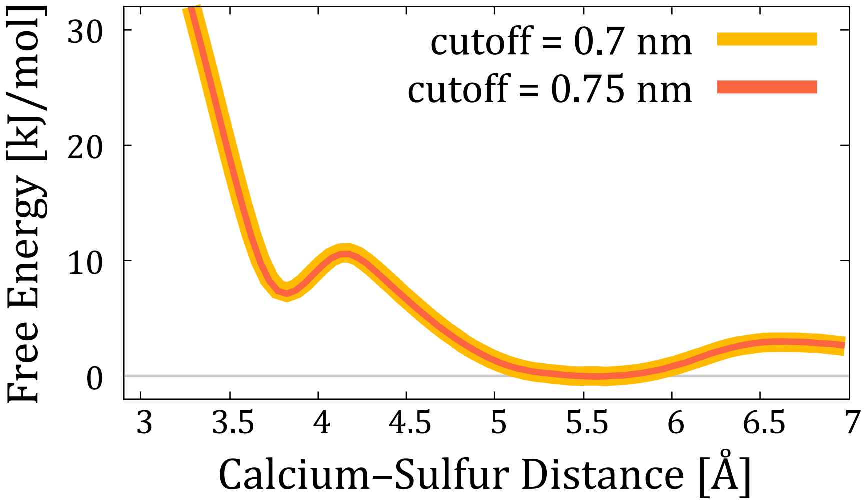


**Figure S15:** Interaction free energy profiles obtained from umbrella sampling simulations of calcium binding to MSO4 using CHARMM36-NBFIX model with cutoff for van der Waals and electrostatic cutoff set to 0.7 nm (orange line, used in all simulations in the main text) or 0.75 nm (red line).

#### S3.9 Role of Water Model on Calcium Binding

A recent work[S42] suggested that different water models as well as using an implicit solvent have various impacts on the conformation of glycosaminoglycan sequences. To check whether water parameterization has an impact on calcium binding to sulfate/sulfamate groups, we run umbrella sampling simulations with GLYCAM06 and GLYCAM-ECC force fields in combination with OPC3 water model[S43] instead of TIP3P. OPC3 water, as well as four-site OPC model,[S44] are often recommended for simulations with AMBER (note that GLYCAM06 is designed to be AMBER-compatible). Our results, Figure S16, show that while OPC3 model brings some changes to the free energy profiles and even somewhat decreases the overestimated preference of monodentate binding in GLYCAM06 simulations, these changes are not substantial to alter the (dis)agreement with the reference AIMD data.

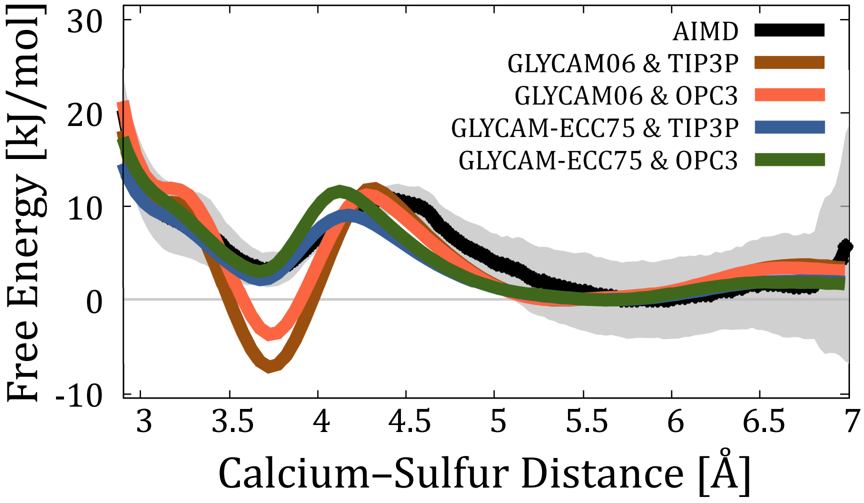


**Figure S16:** Interaction free energy profiles obtained from umbrella sampling simulations of calcium binding to MSO4 using GLYCAM06 or GLYCAM-ECC75 force field with either TIP3P (used in all GLYCAM06 and GLYCAM-ECC simulations in the main text) or OPC3 water model as compared to AIMD data.

The similar conclusions can be made from simulations with CHARMM36-NBFIX and prosECCo75 force fields in combination with TIP4P/2005 water,[S45] Figure S17, which is another highly popular water model often regarded as one of the best rigid nonpolarizable water models.[S46] While the choice of the model has an effect on the free energy profiles (TIP45/2005 promotes the stronger monodenate binding), related changes are not comparable to those associated with the implicit inclusion (charge scaling) of the electronic polarization or even *ad hoc* repulsive interactions within the NBFIX concept.

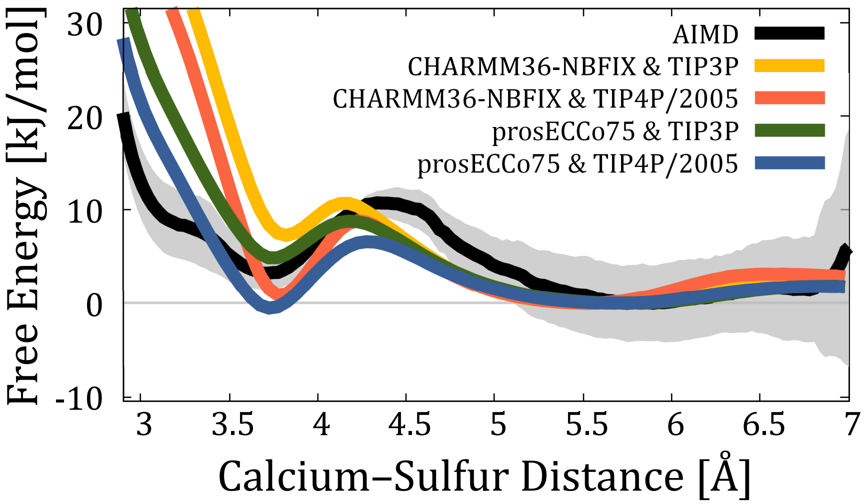


**Figure S17:** Interaction free energy profiles obtained from umbrella sampling simulations of calcium binding to MSO4 using CHARMM36-NBFIX or prosECCo75 force field with either TIP3P (used in all CHARMM36, CHARMM36-NBFIX, and prosECCo75 simulations in the main text) or TIP4P/2005 water model as compared to AIMD data.

#### S3.10 Accelerated Weight Histogram vs. Umbrella Sampling Simulations

To verify if umbrella sampling methodology is suitable for our systems, we also tested accelerated weight histogram (AWH) method to compare the results. Our data, Figure S18, show that both methods provide identical results.

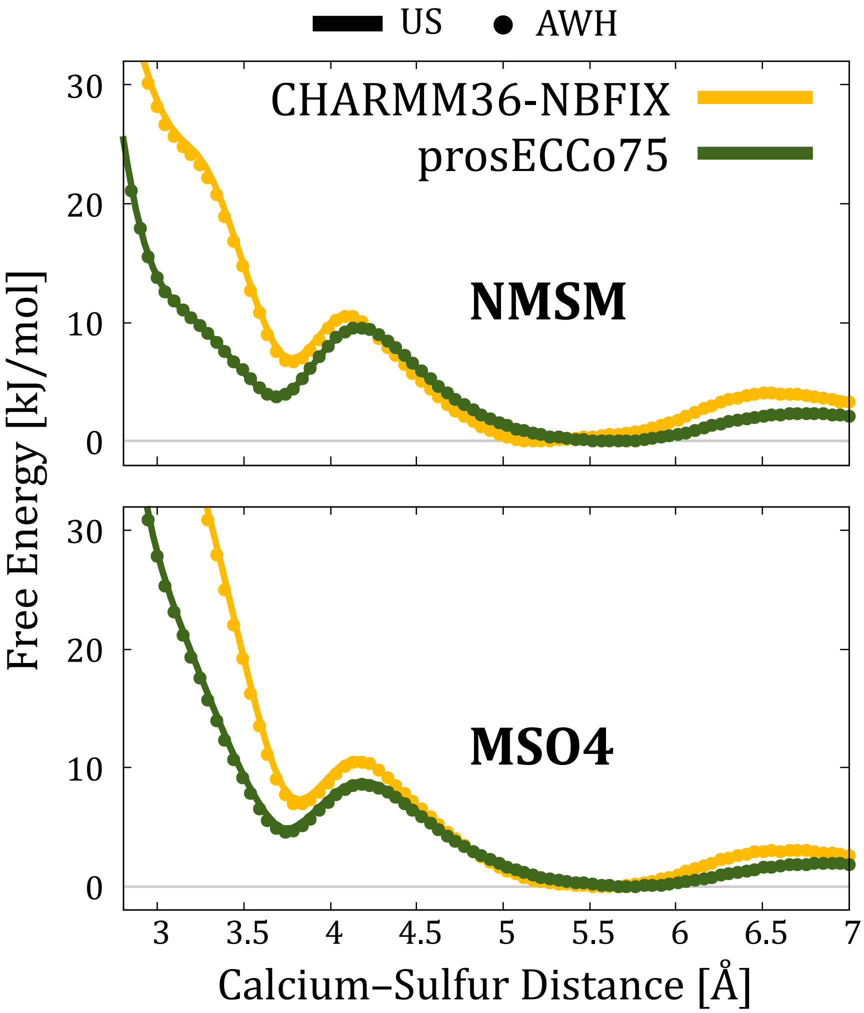


**Figure S18:** Comparison of FFMD interaction free energy profiles obtained from umbrella sampling (US) and accelerated weight histogram (AWH) simulations.

#### S3.11 Additional Free Energy Profiles

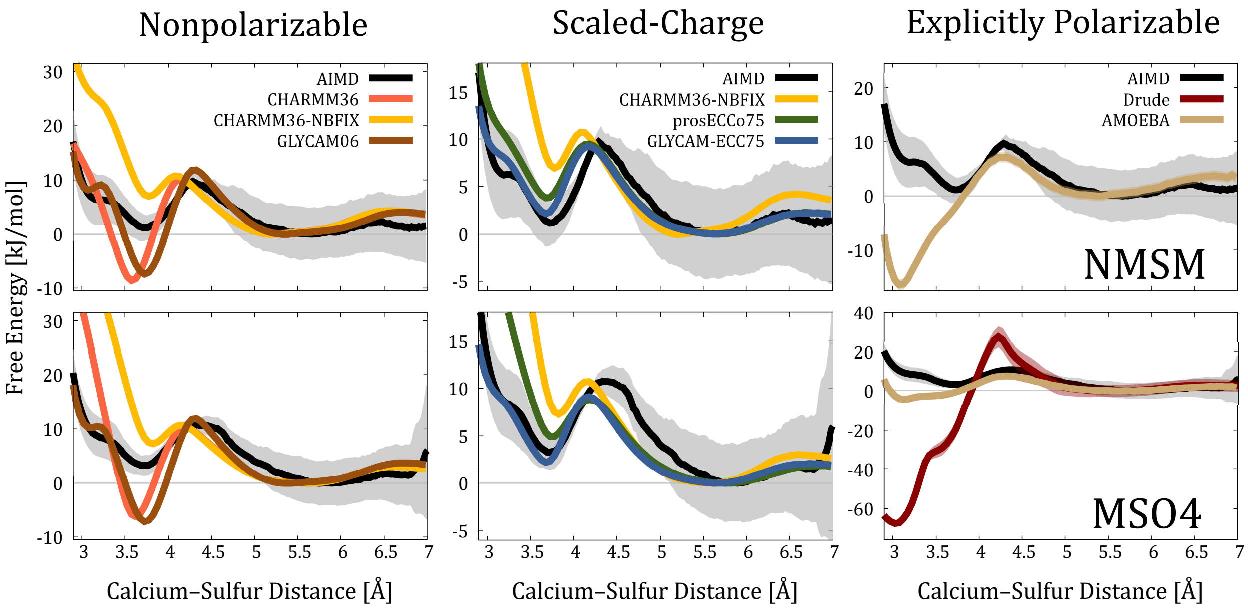


**Figure S19:** The same as Figure 3 but with adjusted vertical scales for better per-force-field visualization.

#### S3.12 Additional Radial Distribution Functions

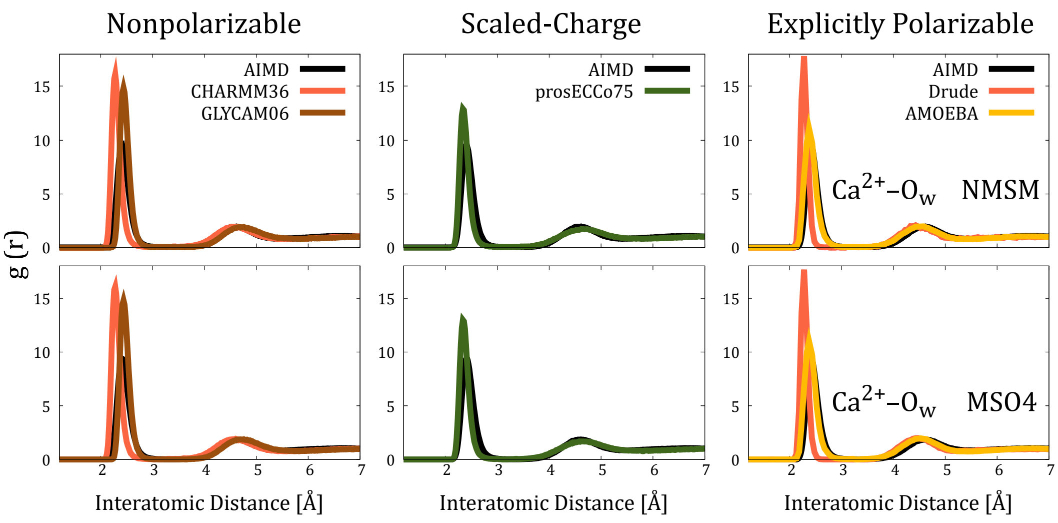


**Figure S20:** $Ca^{2+}$–$O_w$ RDFs collected from AIMD and FFMD simulations.

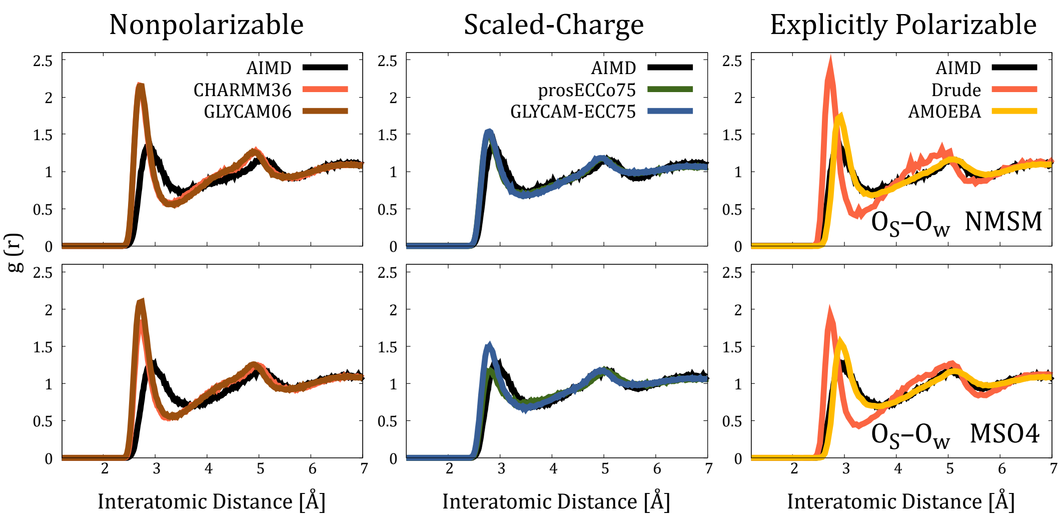


**Figure S21:** $O_S$–$O_w$ RDFss collected from AIMD and FFMD simulations.

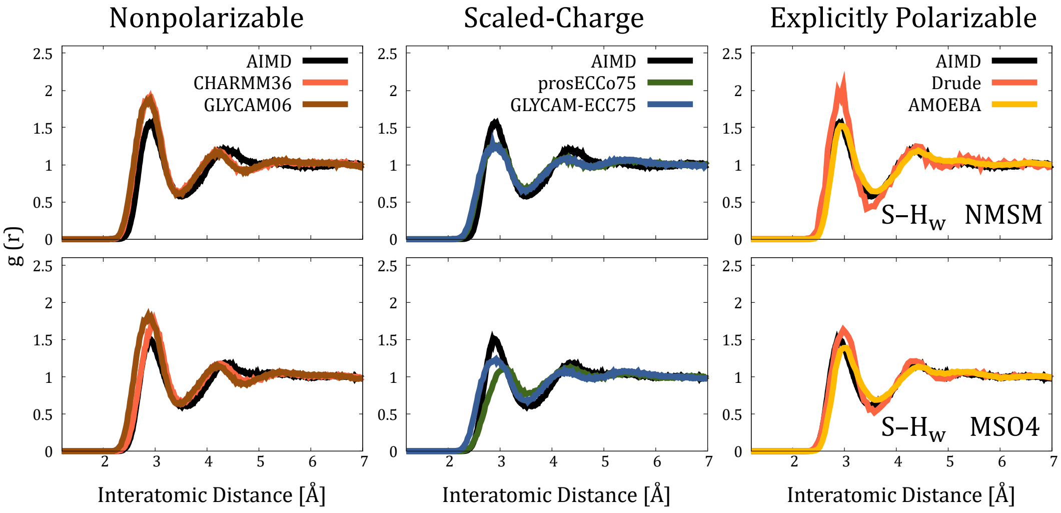


**Figure S22:** S–$H_w$ RDFs collected from AIMD and FFMD simulations.

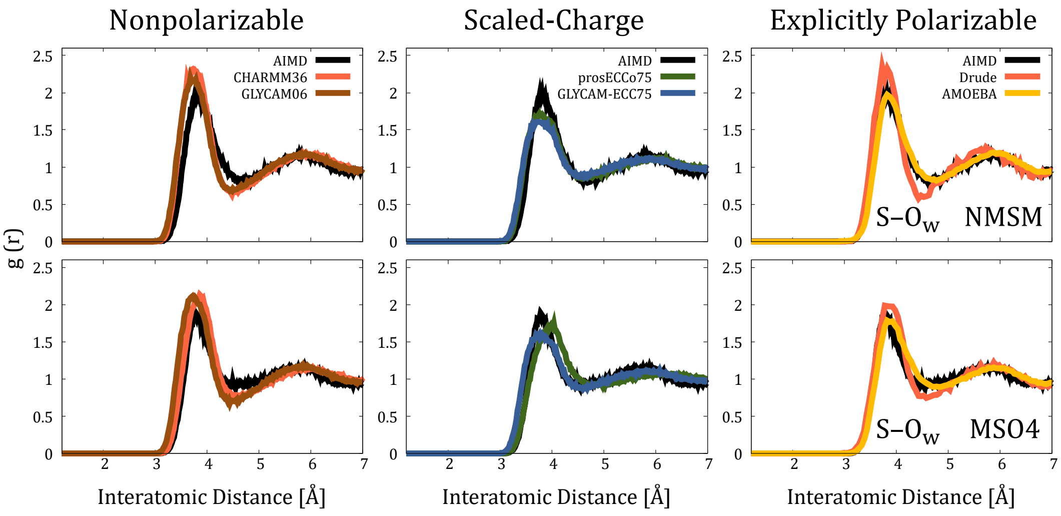


**Figure S23:** S–$O_w$ RDFs collected from AIMD and FFMD simulations.

(S15) MacKerell, A. D. Department of Pharmaceutical Sciences, School of Pharmacy, University of Maryland, Baltimore, Maryland 21201, United States. Personal communication. 2022.

(S16) Lamoureux, G.; Harder, E.; Vorobyov, I. V.; Roux, B.; MacKerell, A. D. A polarizable model of water for molecular dynamics simulations of biomolecules. *Chem. Phys. Lett.* **2006**, *418*, 245–249.

(S17) Yu, H.; Whitfield, T. W.; Harder, E.; Lamoureux, G.; Vorobyov, I.; Anisimov, V. M.; MacKerell, A. D.; Roux, B. Simulating monovalent and divalent ions in aqueous solution using a drude polarizable force field. *J. Chem. Theory Comput.* **2010**, *6*, 774–786.

(S18) Humphrey, W.; Dalke, A.; Schulten, K. VMD: Visual molecular dynamics. *J. Mol. Graph.* **1996**, *14*, 33–38.

(S19) Park, S. J.; Lee, J. J.; Qi, Y.; Kern, N. R.; Lee, H. S.; Jo, S.; Joung, I.; Joo, K.; Lee, J. J.; Im, W. CHARMM-GUI Glycan Modeler for modeling and simulation of carbohydrates and glycoconjugates. *Glycobiology* **2019**, *29*, 320–331.

(S20) Abraham, M. J.; Murtola, T.; Schulz, R.; Páll, S.; Smith, J. C.; Hess, B.; Lindah, E. Gromacs: High performance molecular simulations through multi-level parallelism from laptops to supercomputers. *SoftwareX* **2015**, *1-2*, 19–25.

(S21) Páll, S.; Hess, B. A flexible algorithm for calculating pair interactions on SIMD architectures. *Comput. Phys. Commun.* **2013**, *184*, 2641–2650.

(S22) Nosé, S. A molecular dynamics method for simulations in the canonical ensemble. *Mol. Phys.* **1984**, *52*, 255–268.

(S23) Hoover, W. G. Canonical dynamics: Equilibrium phase-space distributions. *Phys. Rev. A* **1985**, *31*, 1695–1697.

(S24) Parrinello, M.; Rahman, A. Polymorphic transitions in single crystals: A new molecular dynamics method. *J. Appl. Phys.* **1981**, *52*, 7182–7190.

(S25) Essmann, U.; Perera, L.; Berkowitz, M. L.; Darden, T.; Lee, H.; Pedersen, L. G. A smooth particle mesh Ewald method. *J. Chem. Phys.* **1995**, *103*, 8577–8593.

(S26) Steinbach, P. J.; Brooks, B. R. New spherical-cutoff methods for long-range forces in macromolecular simulation. *J. Comput. Chem.* **1994**, *15*, 667–683.

(S27) Hess, B.; Bekker, H.; Berendsen, H. J.; Fraaije, J. G. LINCS: A Linear Constraint Solver for molecular simulations. *J. Comput. Chem.* **1997**, *18*, 1463–1472.

(S28) Hess, B. P-LINCS: A parallel linear constraint solver for molecular simulation. *J. Chem. Theory Comput.* **2008**, *4*, 116–122.

(S29) Jorgensen, W. L.; Chandrasekhar, J.; Madura, J. D.; Impey, R. W.; Klein, M. L. Comparison of simple potential functions for simulating liquid water. *J. Chem. Phys.* **1983**, *79*, 926–935.

(S30) Durell, S. R.; Brooks, B. R.; Ben-Naim, A. Solvent-induced forces between two hydrophilic groups. *J. Phys. Chem.* **1994**, *98*, 2198–2202.

(S31) Miyamoto, S.; Kollman, P. A. Settle: An analytical version of the SHAKE and RATTLE algorithm for rigid water models. *J. Comput. Chem.* **1992**, *13*, 952–962.

(S32) Phillips, J. C.; Hardy, D. J.; Maia, J. D.; Stone, J. E.; Ribeiro, J. V.; Bernardi, R. C.; Buch, R.; Fiorin, G.; Hénin, J.; Jiang, W.; McGreevy, R.; Melo, M. C.; Radak, B. K.; Skeel, R. D.; Singharoy, A.; Wang, Y.; Roux, B.; Aksimentiev, A.; Luthey-Schulten, Z.; Kalé, L. V.; Schulten, K.; Chipot, C.; Tajkhorshid, E. Scalable molecular dynamics on CPU and GPU architectures with NAMD. *J. Chem. Phys.* **2020**, *153*, 44130.

(S33) Fiorin, G.; Klein, M. L.; Hénin, J. Using collective variables to drive molecular dynamics simulations. *Mol. Phys.* **2013**, *111*, 3345–3362.

(S34) Jiang, W.; Hardy, D. J.; Phillips, J. C.; MacKerell, A. D.; Schulten, K.; Roux, B. High-performance scalable molecular dynamics simulations of a polarizable force field based on classical drude oscillators in NAMD. *J. Phys. Chem. Lett.* **2011**, *2*, 87–92.

(S35) Chowdhary, J.; Harder, E.; Lopes, P. E.; Huang, L.; MacKerell, A. D.; Roux, B. A polarizable force field of dipalmitoylphosphatidylcholine based on the classical drude model for molecular dynamics simulations of lipids. *J. Phys. Chem. B* **2013**, *117*, 9142–9160.

(S36) Lemkul, J. A.; Huang, J.; Roux, B.; Mackerell, A. D. An Empirical Polarizable Force Field Based on the Classical Drude Oscillator Model: Development History and Recent Applications. *Chem. Rev.* **2016**, *116*, 4983–5013.

(S37) Tuckerman, M.; Berne, B. J.; Martyna, G. J. Reversible multiple time scale molecular dynamics. *J. Chem. Phys.* **1992**, *97*, 1990–2001.

(S38) Bussi, G.; Donadio, D.; Parrinello, M. Canonical sampling through velocity rescaling. *J. Chem. Phys.* **2007**, *126*, 014101.

(S39) Wu, J. C.; Chattree, G.; Ren, P. Automation of AMOEBA polarizable force field parameterization for small molecules. *Theor. Chem. Acc.* **2012**, *131*, 1–11.

(S40) Kostal, V.; Mason, P. E.; Martinez-Seara, H.; Jungwirth, P. Common Cations Are Not Polarizable: Effects of Dispersion Correction on Hydration Structures from Ab Initio Molecular Dynamics. *J. Phys. Chem. Lett.* **2023**, *14*, 4403–4408.

(S41) Grimme, S.; Antony, J.; Ehrlich, S.; Krieg, H. A consistent and accurate ab initio parametrization of density functional dispersion correction (DFT-D) for the 94 elements H-Pu. *J. Chem. Phys.* **2010**, *132*.

(S42) Marcisz, M.; Samsonov, S. A. Solvent Model Benchmark for Molecular Dynamics of Glycosaminoglycans. *J. Chem. Inf. Model.* **2023**, *63*, 2147–2157.

(S43) Izadi, S.; Onufriev, A. V. Accuracy limit of rigid 3-point water models. *J. Chem. Phys.* **2016**, *145*.

(S44) Izadi, S.; Anandakrishnan, R.; Onufriev, A. V. Building water models: A different approach. *J. Phys. Chem. Lett.* **2014**, *5*, 3863–3871.

(S45) Abascal, J. L.; Vega, C. A general purpose model for the condensed phases of water: TIP4P/2005. *J. Chem. Phys.* **2005**, *123*.

(S46) Vega, C.; Abascal, J. L. Simulating water with rigid non-polarizable models: A general perspective. 2011; `https://pubs.rsc.org/en/content/articlehtml/2011/cp/c1cp22168jhttps://pubs.rsc.org/en/content/articlelanding/2011/cp/c1cp22168j`.
